# Human NMDAR autoantibodies disrupt excitatory-inhibitory balance leading to hippocampal network hypersynchrony

**DOI:** 10.1101/2022.03.04.482796

**Authors:** Mihai Ceanga, Vahid Rahmati, Holger Haselmann, Lars Schmidl, Daniel Hunter, Jakob Kreye, Harald Prüss, Laurent Groc, Stefan Hallermann, Josep Dalmau, Alessandro Ori, Manfred Heckmann, Christian Geis

## Abstract

Specific autoantibodies against the NMDA-receptor (NMDAR) GluN1 subunit cause severe and debilitating NMDAR-encephalitis. Autoantibodies induce prototypic disease symptoms resembling schizophrenia, including psychosis and cognitive dysfunction. Using a mouse passive transfer model applying human monoclonal anti-GluN1-autoantibodies, we observed CA1 pyramidal neuron hypoexcitability, reduced AMPA-receptor (AMPAR) signaling, and faster synaptic inhibition resulting in disrupted excitatory-inhibitory balance. Functional alterations were supported by widespread remodeling of the hippocampal proteome, including changes in glutamatergic and GABAergic neurotransmission. At the network level, anti-GluN1-autoantibodies amplified gamma oscillations and disrupted theta-gamma coupling. A data-informed network model revealed that lower AMPAR strength and faster GABAA-receptor current kinetics chiefly account for these abnormal oscillations. As predicted by our model and evidenced experimentally, positive allosteric modulation of AMPARs alleviated aberrant gamma activity and thus reinforced the causative effects of the excitatory-inhibitory imbalance. Collectively, NMDAR-hypofunction-induced aberrant synaptic, cellular, and network dynamics provide new mechanistic insights into disease symptoms in NMDAR-encephalitis and schizophrenia.

## Introduction

NMDA receptors (NMDARs) are ionotropic glutamate receptors that play a pivotal role in excitatory transmission and synaptic plasticity in the central nervous system (CNS). They are particularly important for memory formation and psychosocial behavior^1^. Hypofunction of NMDARs has been implicated in the pathophysiology of a variety of complex neuropsychiatric disorders, e.g. in dementia or schizophrenia^1, 2^. NMDAR encephalitis, first described in 2007, is a severe autoimmune CNS disorder which is directly and solely caused by highly specific pathogenic autoantibodies (Ab) targeting the aminoterminal domain of the GluN1 subunit of the NMDAR^3, 4^. Affected patients suffer from a prototypic, severe and complex neuropsychiatric syndrome consisting of psychosis, memory dysfunction, delusional thinking, hallucinations, catatonic features, epilepsy, and central hypoventilation^4^. At the molecular level, the binding of Ab to surface-expressed NMDARs induces crosslinking and internalization of these receptors, followed by a reduction in their surface expression in postsynaptic receptor fields^5^. These direct effects of Ab have been investigated in cell culture models and in passive-transfer animal models after intraventricular infusion of patient-derived Ab against GluN1, providing evidence that Ab-induced NMDAR internalization ultimately results in reduction of NMDAR-mediated excitatory postsynaptic currents and diminished NMDAR-dependent long-term potentiation^5, 6^. NMDAR-Ab-induced disease in those models brought about cognitive and behavior disorders reflecting fundamental impairment of brain function. However, it remains largely unknown how the observed molecular changes affect computations at the single-cell and neural network levels.

NMDAR-hypofunction induced through genetic or pharmacological manipulations has been previously used to mimic symptoms of schizophrenia, which resembles key features in the clinical presentation of NMDAR-encephalitis^7, 8^. In models of schizophrenia, NMDAR-hypofunction was found to strongly disturb the synchronized neuronal activity patterns^2, 9, 10^ such as γ-oscillations (ca. 20-90 Hz)^11–13^. Importantly, these high frequency oscillations, through an intricate temporal coordination of the activity of distributed cells or neuronal networks, provide a neural substrate of various cognitive processes and behavior such as memory, learning, and spatial navigation. The aberrant γ-oscillations in schizophrenia are linked to a marked excitatory-inhibitory imbalance due to NMDAR-hypofunction^10^. To uncover the neural and network origins of the NMDAR-Ab-induced cognitive and behavioral abnormalities, we specifically investigate the effect of patient-derived monoclonal antibodies against the NMDAR-GluN1 subunit on the intrinsic cellular properties together with hippocampal proteome remodeling, on the excitatory-inhibitory balance, and on the network oscillations. These investigations may have several important implications. First, they can identify the impaired neural and network coding mechanisms underlying the reported, strictly NMDAR-dependent abnormalities of brain function. Second, they can propose specific therapeutic strategies to rescue these impairments. Third, they can offer further insights into brain dysfunction in other pathologies linked to NMDAR-hypofunction, such as schizophrenia.

## Results

### Hypoexcitability and decreased jitter of synaptically-driven CA1 neuronal output in a mouse model of NMDAR-Ab encephalitis

NMDARs regulate neuronal activity by triggering long-term plasticity mechanisms. Therefore, we hypothesized that Ab-induced NMDAR-hypofunction would affect the neuronal input-output (I/O) function. Here, we specially focused on CA1 pyramidal neurons (CA1-PNs), as the read-outs of hippocampus. The I/O function of CA1-PNs is a fundamental computational property of the neuron, and is mainly governed by the interplay between intrinsic excitability and their synaptic inputs, driven largely by feed-forward excitation (CA3→CA1) and inhibition (CA3→PV→CA1)^14, 15^. To test this hypothesis we used an established passive-transfer mouse model with chronic intraventricular delivery^6^. We applied human monoclonal antibodies against the aminoterminal domain of NMDAR-GluN1 subunit derived from antibody-secreting cells of cerebrospinal fluid of patients with NMDAR encephalitis^16^ or monoclonal control Ab (Fig. 1a). We then measured the synaptically-driven I/O function of CA1-PNs in acute hippocampal slices after incremental stimulation of the Schaffer collaterals (SC) (Fig. 1b,c,d). NMDAR-Ab treatment led to an increase in the I/O threshold (EPSP slope with a 50% action potential, AP, generation probability), but did not change the I/O gain (Fig. 1d,e,f,g). Since changes in the excitatory-inhibitory balance (E-I-balance) affect the gain and threshold of the I/O function^14^ as well as AP timing^17, 18^, we next investigated subthreshold responses and AP properties of CA1-PNs. After NMDAR-Ab treatment, EPSP amplitudes were smaller for the same rise slope (Fig. 1h) despite an unchanged membrane time constant (Supplementary Fig. 1), suggesting an alteration in the E-I-balance. We observed an effective loss of low-amplitude EPSPs, a deviation from the expected log-normal distribution^19^, and a marked reduction of EPSP latency dispersion in favor of shorter latencies (Fig. 1i). Consequently, AP-eliciting EPSPs had higher slopes, while AP-latency distribution was narrower (Fig. 1j). Furthermore, EPSP and AP jitter were lower after NMDAR-Ab treatment (Fig. 1k, l). Of note, the resting membrane potential (Vrest) and other AP properties, including AP threshold, remained unchanged (Supplementary Fig. 1). In conclusion, NMDAR-Ab induces synaptic hypoexcitability of CA1-PNs, resulting in an increased I/O threshold, as well as a decreased EPSP and AP jitter.

**Figure 1.**
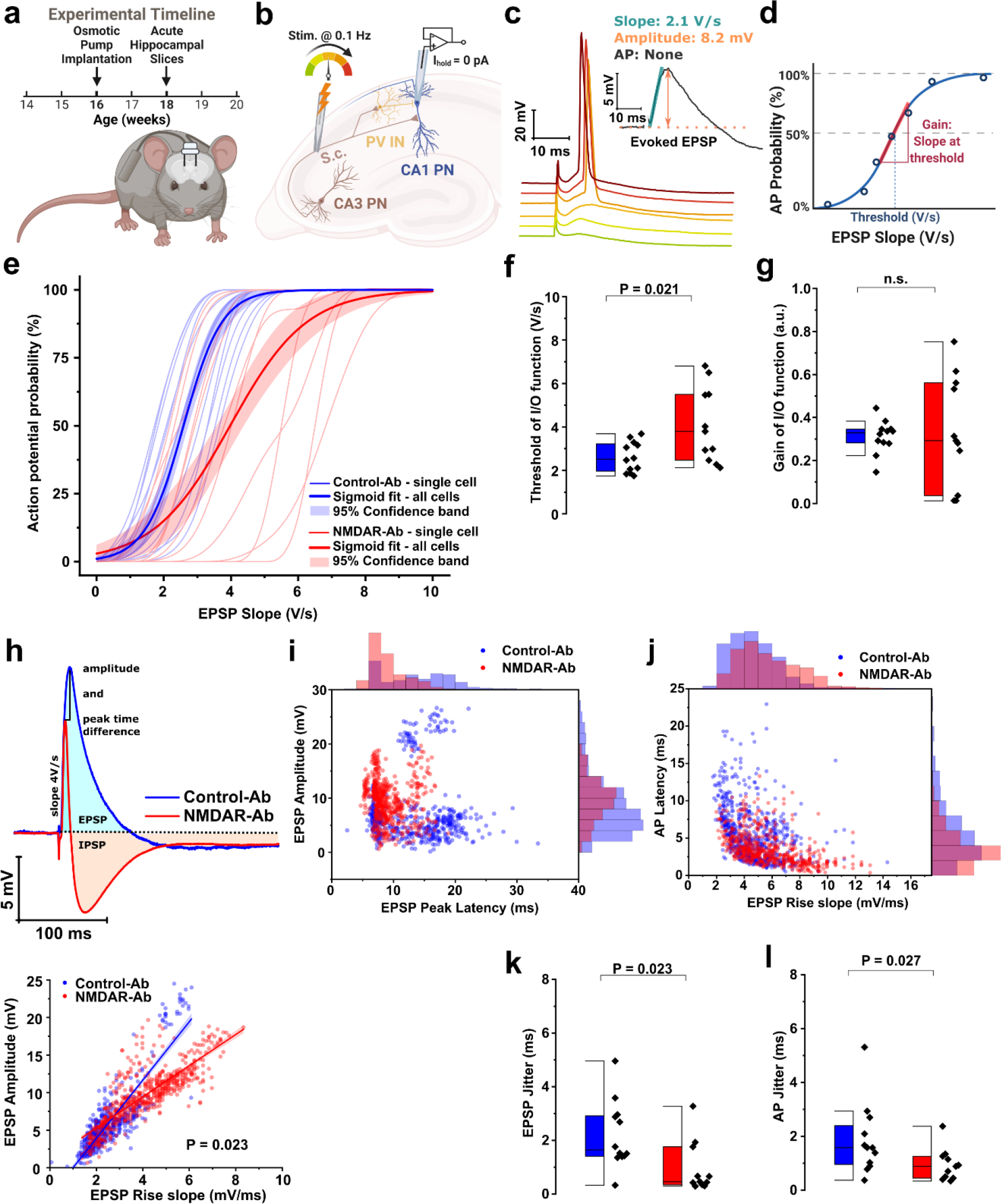
NMDAR-Ab increases the I/O threshold and reduces EPSP and AP jitter in CA1 pyramidal neurons (PNs). **(a)** Schematic timeline of the experiments: sixteen-week-old male mice were implanted with biventricular catheters connected to osmotic pumps, whereby receiving human monoclonal NMDAR-Ab continuously for 2 weeks, after which acute hippocampal slices were prepared. **(b)** The cellular excitability of the feed-forward hippocampal microcircuit was investigated using current-clamp recordings of CA1-PN at a holding current (Ihold) of 0 pA, followed by incremental stimulation of the Schaffer collaterals (SC). **(c)** Example responses to incremental stimulation intensities, showing an increase in the slope and amplitude of the EPSP, which is followed by a delayed IPSP or an AP. The time latency of AP is reduced at higher stimulus intensities. *Inset:* the slope and amplitude of subthreshold EPSPs, as well as the presence of an AP were measured. **(d)** The schematic of I/O function, plotted as the binned EPSP slope versus the AP probability per bin, takes a sigmoid form characterized by a threshold (EPSP slope with 50% AP probability) and a gain (slope of the I/O curve at threshold). **(e)** NMDAR-Ab shifts the I/O function of CA1-PN to the right, indicating hypoexcitability. Thin lines are single cell sigmoid fits of I/O data. Solid lines and shaded areas depict the mean and 95% CI bands of fits per treatment group. **(f)** The NMDAR-Ab increases the threshold of I/O curves. **(g)** The group-averaged gain of I/O curves. **(h)** *Upper panel:* Example biphasic responses (EPSP+IPSP) in Control-Ab and NMDAR-Ab groups, showing the same EPSP slope but different EPSP amplitudes, peak latencies, and overall morphologies. *Lower panel:* Scatter plot of all-cells’ EPSP events with the linear fits showing a less increase in EPSP amplitude at steeper EPSP slopes under NMDAR-Ab. **(i)** Scatter plot and marginal histograms of latency and amplitude distribution of all-cells’ subthreshold EPSP events. NMDAR-Ab reduced frequencies of both long-latency and low-amplitude EPSP events, and made EPSP amplitude distribution nearly symmetric. **(j)** Same as (i), but for EPSP slope and AP latency of all individual AP-eliciting EPSP responses. The distribution of AP-eliciting EPSP slopes exhibits less positive-skewness after NMDAR-Ab treatment, rendering the AP-generation by low EPSP slopes less likely. In contrast, AP latencies exhibit more positive-skewness, indicating confined timing for AP generation. **(k)** NMDAR-Ab decreases the EPSP jitter, calculated as the SD of EPSP latency at I/O threshold. **(l)** NMDAR-Ab decreases AP jitter, calculated as the SD of the AP latency at 2x the I/O threshold slope (or at 10V/s). The details of all statistical tests and sample sizes of data used throughout the figures and main text can be found in Supplementary Table 1.

### Reduced AP firing after NMDAR-Ab treatment

Since intrinsic-excitability plasticity often co-occurs with synaptic plasticity^15^, we next hypothesized that NMDAR-Ab would alter the passive and active excitability characteristics of CA1-PNs. As these neurons show a prominent resonance in the theta band (4-12 Hz) that can promote their information transfer efficiency during subthreshold and network oscillations ^20^, we also measured membrane impedance. NMDAR-Ab caused a reduction in both impedance and input resistance (Rin), with no change in the resonant frequency, Vrest, or membrane tau (Fig. 2a-d, Supplementary Fig. 1). Upon current-step injections, the mean instantaneous frequency was decreased by NMDAR-Ab, but we did not find a change in the number of elicited APs or in the spike frequency adaptation (Fig. 2e-h). Of note, APs in CA1-PNs can occur not only after depolarization, but also as post-inhibitory rebound spikes. Upon membrane hyperpolarization steps, a lower proportion of PNs exhibited rebound APs in NMDAR-Ab group (Fig. 2i). Membrane voltage sag and properties of APs induced by the current injections at the soma remained unchanged, similarly to our synaptic stimulation data (Supplementary Fig. 1). In summary, NMDAR-Ab downregulates CA1-PNs’ excitability by reducing membrane impedance and AP frequency, and restricting rebound-AP generation.

**Figure 2.**
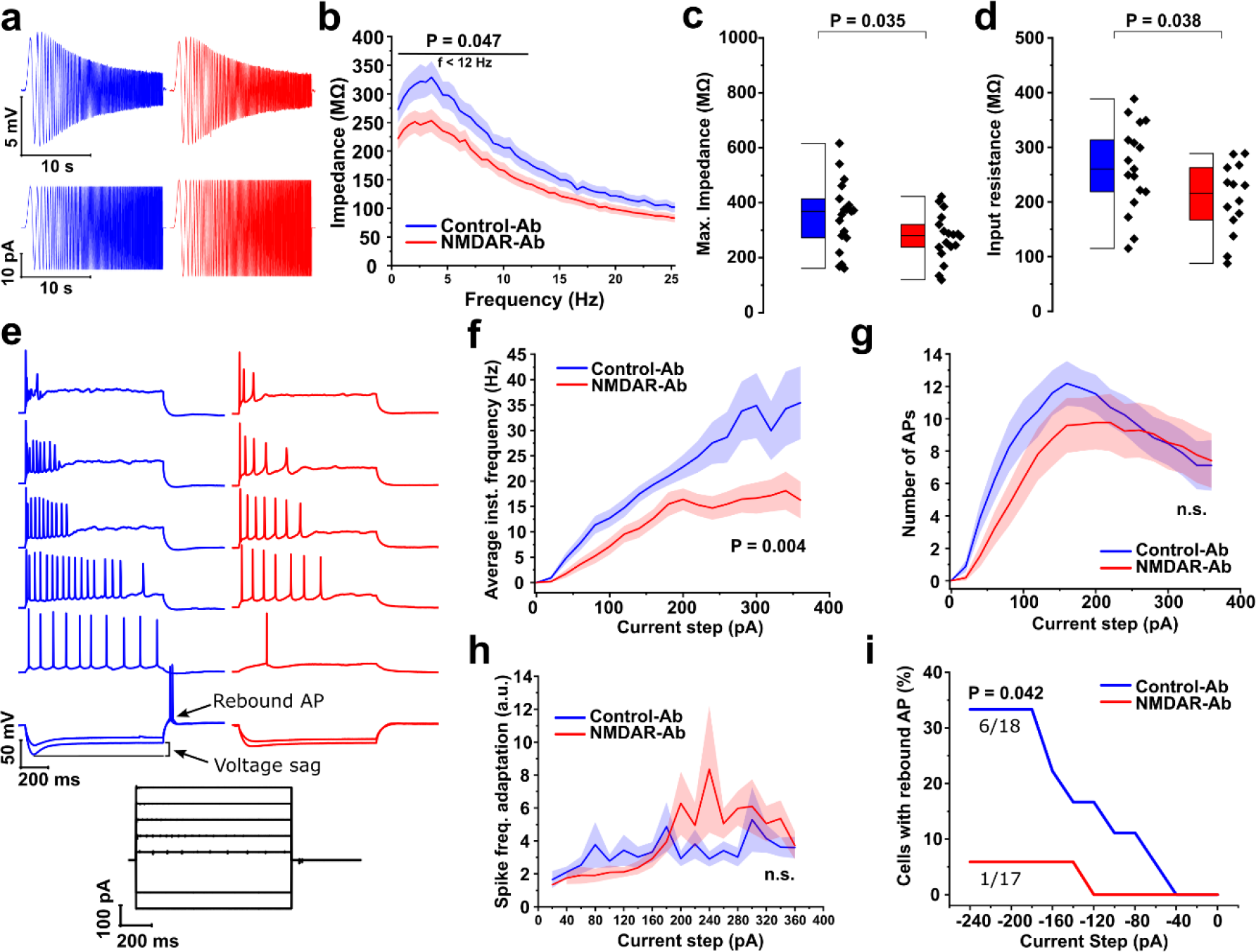
NMDAR-Ab induces intrinsic hypoexcitability of CA1 pyramidal neurons (PNs). **(a)** Example traces of the membrane impedance showing the subthreshold resonance of CA1-PN (upper traces) during injection of a chirp current with linearly increasing frequency (lower traces). **(b)** The impedance drops after NMDAR-Ab treatment for frequencies below 12 Hz. The mean (solid line) ± SEM (shaded area) **(c)** The maximum impedance, calculated at peak (i.e. resonant) frequency, decreases after NMDAR-Ab treatment. **(d)** The input resistance decreases after NMDAR-Ab treatment. **(e)** Example response traces during 1-second current injection at the soma (-250pA to 360pA in 20pA steps, from 0pA holding current), showing the induction of AP trains, or voltage sag at negative membrane potentials followed by rebound APs. Note the reduced firing propensity in NMDAR-Ab group. **(f)** The average instantaneous firing frequency of CA1-PNs drops after NMDAR-Ab treatment. **(g)** The number of APs per current step (1-second) and **(h)** the spike frequency adaptation (ISIlast/ISIfirst) remained almost unchanged across the treatments. **(i)** Rebound APs were nearly abolished by NMDAR-Ab (at Icmd = -240pA). See Supplementary Table 1 for statistical details.

### Reduced AMPA strength and faster GABAA kinetics in CA1 pyramidal neurons after NMDAR-Ab treatment

While membrane impedance affects EPSP and IPSP proportionately, an increased I/O threshold with unchanged gain suggests reduced synaptic excitation^14^. We therefore investigated the underlying synaptic currents by measuring spontaneous excitatory (sEPSC, AMPA-mediated) and inhibitory (sIPSC, GABAA-mediated) inputs at the respective reversal potentials of inhibition and excitation (Fig. 3, Supplementary Fig. 2a). NMDAR-Ab reduced the amplitude, decay time constant (τ_*d*_) and total charge transfer of sEPSC onto CA1-PNs, but did not affect event frequency (Fig. 3a-g). For sIPSC, we observed no effect of NMDAR-Ab on amplitude and frequency, but a pronounced decrease in τ_*d*_ and charge transfer (Fig. 3h-n, Supplementary Fig. 2b). Furthermore, the peak of sEPSC, but not sIPSC, frequency distribution was misaligned with the resonant frequency of CA1-PNs, and sEPSC event density was lower at the resonant frequency, thereby possibly contributing to reduced amplification of EPSPs specifically (Supplementary Fig. 2c,d). Additionally, by analyzing dendritic and spine structure in CA1-PNs, we investigated the structural basis of our observed changes in synaptic conductances. Reconstructing CA1-PNs revealed however almost no effect of NMDAR-Ab on dendritic structures and spine distributions (Supplementary Fig. 3). Thus, functional rather than structural changes underlie the observed synaptic alterations. These data demonstrate decreased amplitude of AMPAR-mediated currents and faster decaying kinetics of GABAAR-mediated currents after NMDAR-Ab treatment.

**Figure 3.**
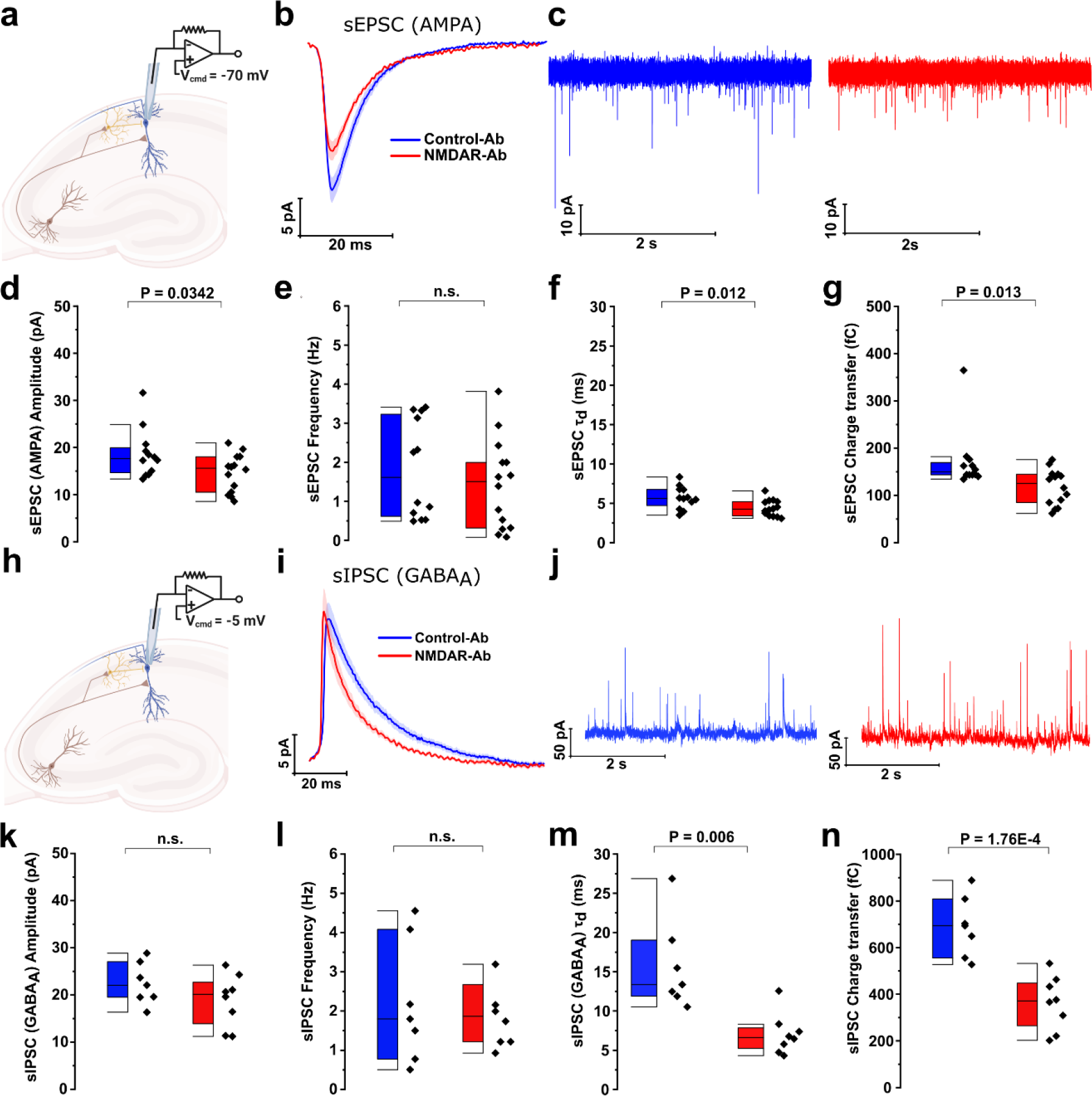
NMDAR-Ab differentially affects AMPAR-and GABAAR-mediated inputs onto CA1 pyramidal neurons (PNs). **(a)** CA1-PNs were held at *E*Inhibition (-70mV) to record spontaneous excitatory postsynaptic AMPAR currents (sEPSC). **(b)** Average sEPSC shape, showing the reduced sEPSC amplitude under NMDAR-Ab. The mean (solid line) ± SEM (shaded area). **(c)** Example recordings of sEPSC from CA1-PNs. **(d-g)** NMDAR-Ab reduced (d) the amplitude, (f) decay time constant, and (g) charge transfer, but not (e) frequency of sEPSC. **(h)** CA1-PNs were held at *E*Excitation (-5mV) to record spontaneous inhibitory GABAAR postsynaptic currents (sIPSC). **(i-n)** Same as (b-g), but for sIPSC. Note, under NMDAR-Ab, neither (k) the amplitude nor (l) the frequency of sIPSC was reduced, but there was a decrease in (m) the sIPSC decay time constant and (n) charge transfer of sIPSC. See Supplementary Table 1 for statistical details.

### NMDAR-Ab induces synaptic E-I-imbalance and cell-specific changes in short-term plasticity

Because spontaneous synaptic currents reflect global, largely uncorrelated input onto CA1-PN, we next focused on coordinated evoked events after stimulating the CA3-to-CA1 feed-forward microcircuit and measured the effect of NMDAR-Ab on E-I-balance. We recorded CA1-PNs’ compound postsynaptic currents at an intermediate holding potential (-35mV, range -40mV to -25mV) in response to single-pulse supramaximal SC stimulation. Evoked responses comprise an initial monosynaptic excitatory component (eESPC), followed by a delayed bisynaptic inhibitory component (eIPSC) mediated by parvalbumin-positive interneurons (PV-INs)^21^ (Fig. 4a,b). Corroborating our aforementioned results, NMDAR-Ab increased the inhibitory-excitatory ratio (IER), indicating a shift towards stronger inhibition relative to excitation (Fig. 4b,c). The integration window (the temporal distance between the peaks of eEPSC and eIPSC) was unchanged (Fig. 4d). To disentangle the individual contributions of excitation and inhibition, we again stimulated SC at the reversal potentials of inhibition (-70mV) and excitation (-5mV) and recorded monosynaptic eEPSC and bisynaptic eIPSC, respectively. The I/O curve and maximal amplitudes of eEPSC were reduced after NMDAR-Ab treatment, with no evidence for a change in τ_*d*_ (Fig. 4e-h). Conversely, we confirmed both the unchanged amplitude and faster decay kinetics of bisynaptic eIPSC (Fig. 4i-l), as in our sIPSC data (Fig. 3h-n).

**Figure 4.**
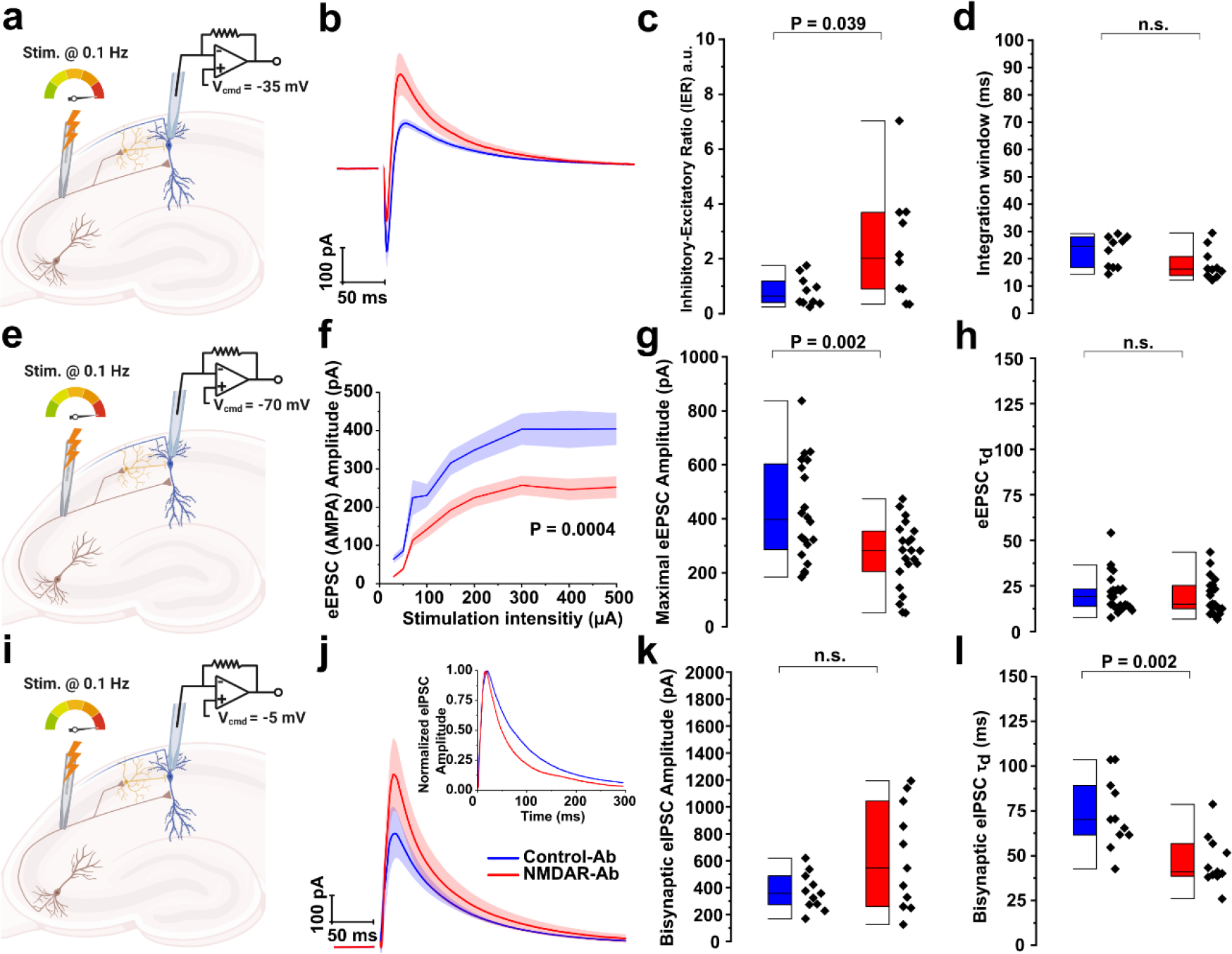
NMDAR-Ab induces E-I-imbalance in CA1 feed-forward inhibition circuit through reduced excitation. **(a-d)** Evoked biphasic postsynaptic response. CA1-PNs were held at an intermediary holding potential (Vcmd) of -35 mV during supramaximal stimulation of Schaffer collaterals (SC). **(a)** Schematic of recording protocol. **(b)** Average biphasic response composed of an initial monosynaptic excitatory component (eEPSC) preceding a delayed, bisynaptic inhibitory component (eIPSC). The mean (solid line) ± SEM (shaded area). **(c)** The inhibitory-excitatory ratio (IER) was increased after NMDAR-Ab treatment. **(d)** The integration window (time interval between peaks of excitation and inhibition) remained unchanged. **(e-h)** Evoked monosynaptic excitatory responses (eEPSC). CA1-PNs were held at a Vcmd of -70 mV during single-pulse incremental SC stimulation. **(e)** Schematic of recording protocol. **(f)** The I/O curve of eEPSC amplitude was shifted downward after NMDAR-Ab treatment. **(g)** The maximal eEPSC amplitude, i.e. the plateau level in (f), was reduced by NMDAR-Ab. **(h)** There was no change in the decay time-constant of eEPSC across the treatments. **(i-l)** Evoked bisynaptic inhibitory responses (eIPSC). CA1-PNs were held at a Vcmd of -5 mV during supramaximal stimulation of SC. **(i)** Schematic of recording protocol. **(j)** Average eIPSC onto CA1-PNs. *Inset:* Average normalized eIPSCs (scaled to have same amplitudes) traces show the faster decay kinetics of eIPSC after NMDAR-Ab treatment. **(k)** NMDAR-Ab did not change the eIPSC amplitude but **(l)** reduced the eIPSC decay time-constant.

A recent study found changes in serial dependence in patients with NMDAR-encephalitis^22^, and computational modeling has linked serial dependence with short-term synaptic plasticity (STP)^23, 24^. Furthermore, IER contains intricate STP dynamics which can enhance spiking probability and reduce jitter in CA1^25^. We therefore evaluated the effect of NMDAR-Abs on STP. Under Control-Ab, corroborating previous results^25^, IER decreased and the integration time-window increased on the second pulse, hence promoting excitation and prolonging the time for integration of excitatory inputs (Supplementary Fig. 4a-c). By contrast, under NMDAR-Ab, IER on the second pulse remained higher, and the integration window remained narrow, indicating less excitation during STP (Supplementary Fig. 4a-c). Biophysically, this shift can arise from a decreased facilitation of CA3→CA1-PN synapses or an attenuated depression at PV→CA1-PN synapses (further PV interneuron recruitment through facilitation of CA3→PV is unlikely since stimulation was supramaximal). We therefore measured STP of eEPSC and found no difference in the facilitatory response under NMDAR-Ab (Supplementary Fig. 4d,e). Instead, we found a pronounced NMDAR-Ab-induced attenuation of the short-term depression in eIPSC (Supplementary Fig. 4f,g). Finally, we investigated whether excitatory input is affected upon high-frequency stimulation (100 pulses at 20Hz, Supplementary Fig. 4h). Normalized eEPSC amplitudes decreased progressively and reached lower steady-state values under NMDAR-Ab (Supplementary Fig. 4i-k), indicating persisting suppression of excitation during high-frequency stimulation.

In conclusion, NMDAR-Abs impair the hippocampal feed-forward microcircuit by altering the synaptic E-I-balance through reduced excitation and faster decay kinetics of inhibition.

### Proteome remodeling underlies the functional changes induced by NMDAR-Ab

Since NMDARs are centrally involved in synaptic plasticity, we hypothesized that NMDAR-Ab directly or indirectly alter hippocampal protein expression. We analyzed protein abundance changes in NMDAR-Ab-treated versus Control-Ab-treated hippocampi using Data Independent Acquisition mass spectrometry. We found significant abundance changes for 667 out of 4938 identified protein groups (Fig. 5a, Supplementary Online Data). To obtain mechanistic insight into affected pathways, we performed an over-representation analysis using KEGG annotation. The long-term potentiation, glutamatergic synapse and calcium signaling pathways were identified among the significantly altered pathways by NMDAR-Ab (Fig. 5b). We next conducted a gene ontology (GO) over-representation analysis that showed a specific and significant regulation of pathways associated with synaptic transmission, second-messenger mediated signaling, synapse organization, and membrane excitability, among others (Fig. 5c). Accounting for the known functional effects of NMDAR-Ab, the levels of Grin1, Grin2a and Grin2b decreased (Fig. 5d,e). However, as suspected from our functional data, the abundance of ionotropic and metabotropic glutamate receptors was also broadly affected, including AMPAR (Gria1, Gria2), kainate receptors (KAR, Grik3), and group I (Grm1, Grm5), II (Grm2, Grm3) and III (Grm 7) metabotropic glutamate receptors (Fig. 5a,d,e Supplementary Online Data). In particular, our findings of reduced EPSC peak amplitudes and faster IPSC dynamics were corroborated by a reduction in the protein levels of Gria1 and Gria2, as well as Gabra5, reflecting α5 subunit containing GABAA receptors, which form dendritically localized receptors with slow kinetics and are preferentially activated by somatostatin interneurons^26^ (Fig. 5d,e). Overall, NMDAR-Ab induced proteome remodeling affects several fundamental neuronal biochemical pathways and proteins, including NMDAR, AMPAR, and GABAAR-alpha5-subunits, as molecular correlates of our observed electrophysiological changes.

**Figure 5.**
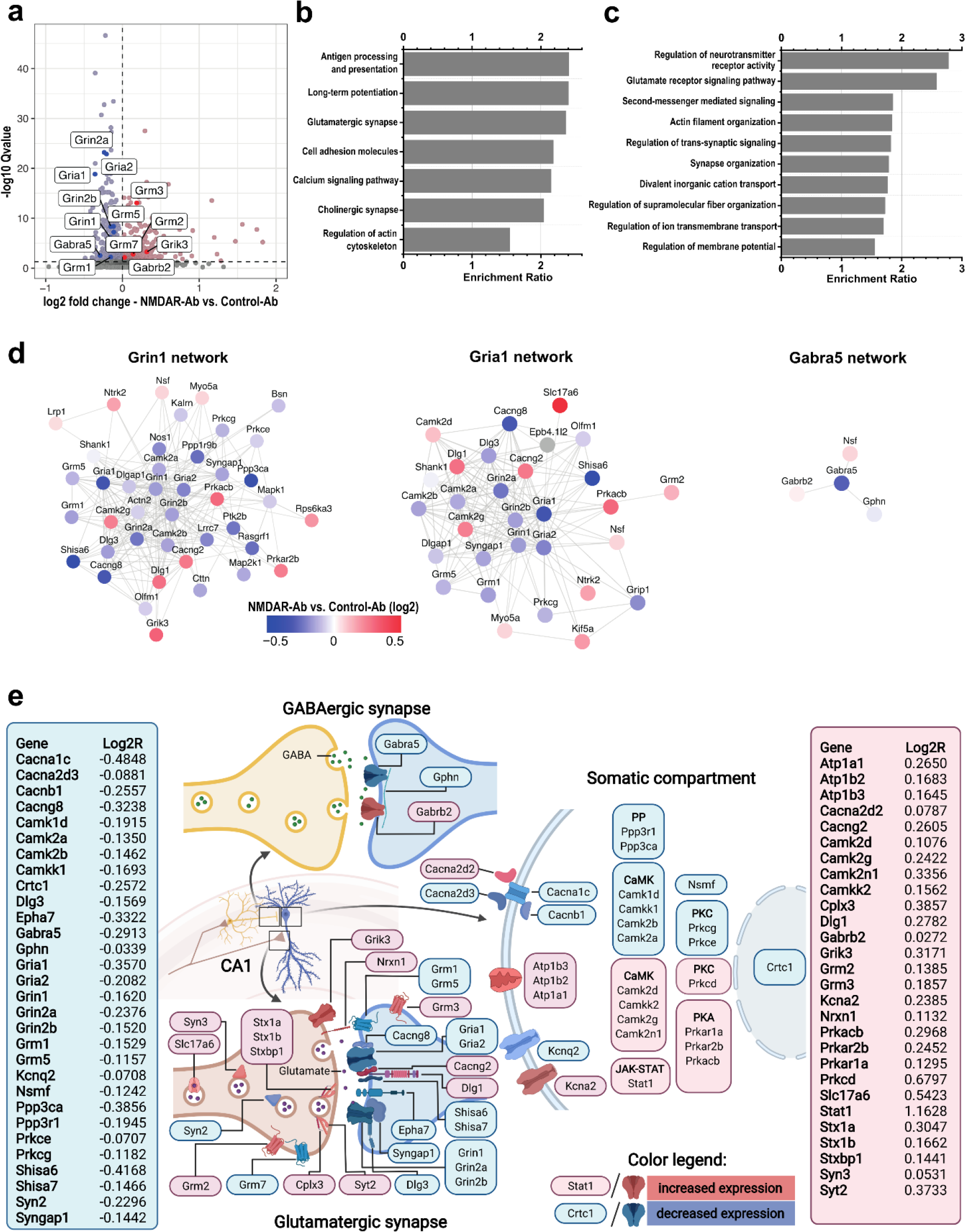
NMDAR-Ab induced proteome changes. **(a)** Volcano plot of quantified proteins in hippocampi following NMDAR-Ab or Control-Ab treatment: n=10 mice for each experimental group. Proteins that significantly (q<0.05) increased or decreased abundance following NMDAR-Ab treatment are highlighted in red and blue, respectively. Glutamate receptors affected by treatment with NMDAR-Ab are annotated. **(b,c)** KEGG pathways (b) and GO biological processes (c) over-represented (FDR<0.05) among proteins affected by NMDAR-Ab treatment. **(d)** Network analysis centered on Grin1 *(left)*, Gria1 *(middle)* and Gabra5 *(right)* showing interacting proteins affected by NMDAR-Ab treatment. Protein-protein interactions were retrieved from STRING^71^ using a high confidence score (>=0.7). **(e)** Graphic representation of the more relevant proteomic changes following NMDAR-Ab treatment, highlighting synapse-, membrane-and enzyme/plasticity-related protein changes (see Supplementary Online Data for the complete list of quantified proteins).

### Abnormal amplification of γ-oscillations by NMDAR-Ab

Excitatory and inhibitory signaling onto PN and IN (majorly the perisomatic-mediated inhibition by e.g. PV-INs) determine γ-oscillation (20-90 Hz) dynamics^2, 27–29^. As γ-oscillations have key roles in various global and hippocampal functions such as attentional selection, encoding, and retrieval of memory traces, among others^29^, we hypothesized that NMDAR-Ab alters γ-oscillations in the hippocampal network.

To test this hypothesis, we used high frequency stimulation (HFS) of SC in acute hippocampal slices of mice treated with NMDAR-Ab or Control-Ab and recorded the extracellular local field potential (LFP) in CA1 stratum pyramidale. HFS induced transient high frequency oscillations in both groups (Fig. 6a). These oscillations were stronger under NMDAR-Ab (Fig. 6b-e) and were mainly confined to the γ-band (30-90Hz) in both groups (Fig. 6c, 6e-inset). In contrast to power strength, there was no change in their peak frequency and bandwidth (Fig. 6f, 6e-inset).

**Figure 6.**
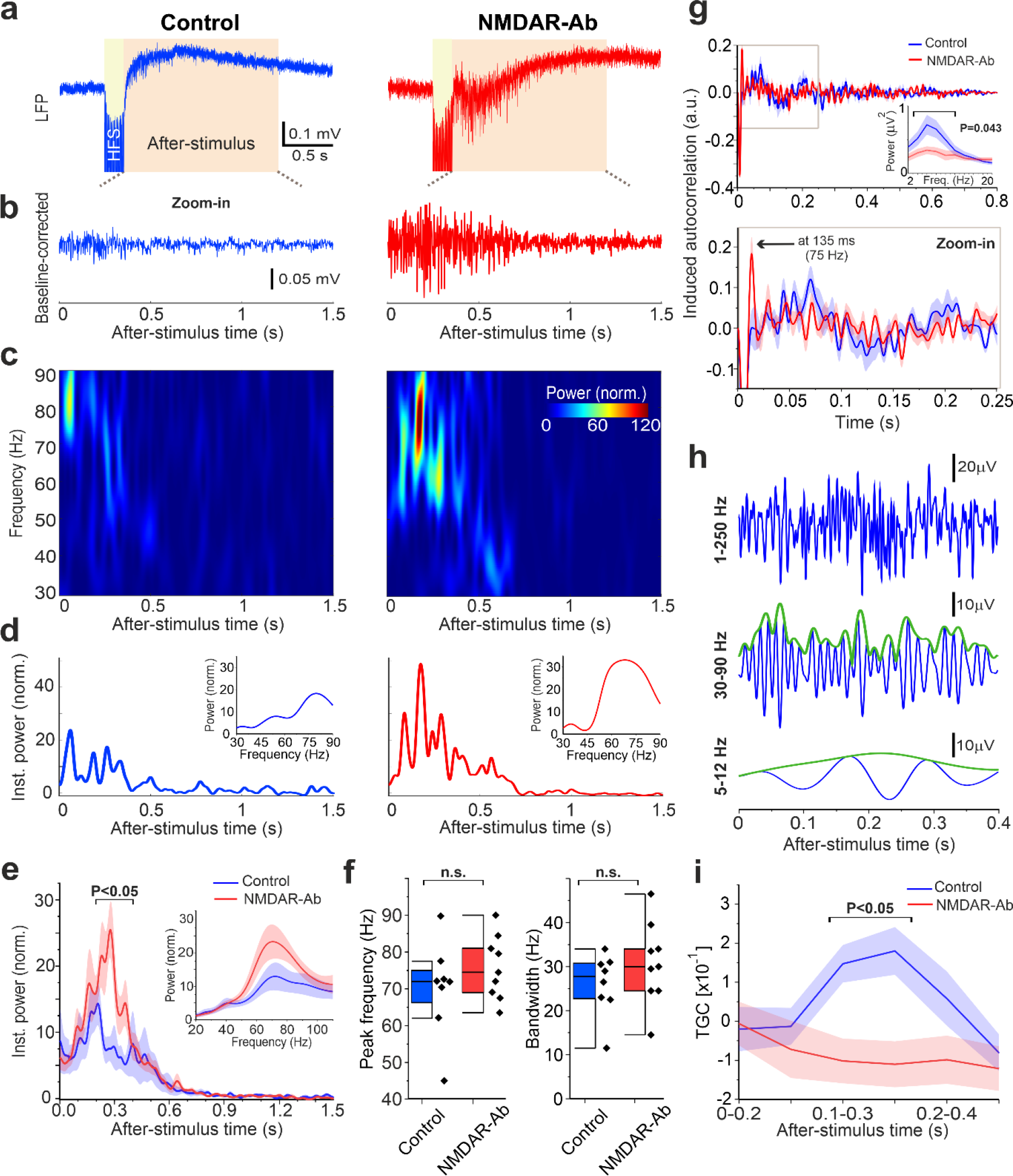
NMDAR-Ab amplifies the transiently induced γ-oscillations in CA1. **(a)** Example ∼3-seconds signals (< 500 Hz) of 10-seconds extracellular local field potential (LFP) recordings of induced network oscillations, from CA1 hippocampal slices, using high frequency stimulation (HFS, 20 pulses at 100Hz with supramaximal stimulation). Note NMDAR-Ab increases the amplitude of induced high frequency oscillations **(b)** Zoom-in of 1.5-seconds of baseline-corrected LFPs (1.1-250 Hz) of those shown in (a) after the HFS. **(c)** The time-frequency plots of power in the LFPs shown in (b), for gamma band (30-90 Hz). Color encodes the power of the after-HFS signal relative to that of the baseline (i.e. before-HFS signal). **(d)** Same as (c), but for the total instantaneous γ-oscillations power, computed as mean power at each time point of the matrices in (c). *Inset*: Wavelet-based power spectrum of HFS-induced γ-oscillations, computed as the mean power at each frequency level of the matrices in (c), over the first 400 milliseconds. **(e)** HFS induces abnormal, stronger γ-oscillations under NMDAR-Ab. Same as (d), but for all recorded slices in the two groups: n=8 for Control, n=9 for NMDAR-Ab. *Inset*: Same as the insets in (d), but for all recorded slices. An extended frequency range is used for better visualization. The mean (solid line) ± SEM (shaded area). (**f**) NMDAR-Ab has no effect on the peak frequency and bandwidth (at half peak-power) of the induced γ-oscillations. **(g)** θ-oscillations are virtually disrupted by NMDAR-Ab. *Top*: Autocorrelograms of the baseline-corrected LFPs after the stimulus offset. *Inset*: Multi-taper power spectrum of the LFPs in theta-band, showing a reduction in the induced θ-oscillatory power under NMDAR-Ab. *Bottom*: Zoom-in of the gray square in the upper panel. **(h)** *Top*: The first 400-milliseconds of the Control LFP in (b). *Middle*: The same signal but band-pass filtered in the gamma band (blue), overlaid by its amplitude envelope (green). *Bottom*: Same as *Middle*, but for the theta band. **(i)** Theta-gamma-comodulation (TGC) is reduced by NMDAR-Ab. TGC was computed for each window of size 0.2 s, with a sliding step of 0.050 s. See Supplementary Table 1 for statistical details.

Furthermore, HFS also induced some concurrent, relatively weak θ-oscillations (5-12Hz) (Fig. 6g,h). Hippocampal θ-nested γ-oscillations are thought to represent a fundamental neural communication mechanism^30, 31^. θ-oscillations were seemingly disturbed and became less contingent with γ-oscillations under NMDAR-Ab (Fig. 6g). The Theta-Gamma-Comodulation (TGC, the Pearson correlation between the amplitude envelop of each LFP signal in theta- and gamma-band, see Methods) was reduced under NMDAR-Ab, particularly between 100-350ms (Fig. 6h,i), which mainly relates to the period of significant increase in the power of γ-oscillations (Fig. 6e). Taken together, these results indicate that NMDAR-Ab amplifies the transiently induced γ-oscillations and, at the same time, tends to impair CA1 information transfer mechanism of the theta-nested γ-oscillations.

### A CA1 neural network model suggests disinhibition of the PN-PV^+^ subnetwork as key to abnormal γ-oscillation amplification

Our experimental data revealed an amplification of γ-oscillations due to NMDAR-Ab, despite synaptic and intrinsic CA1-PN hypoexcitability. To gain mechanistic insights into this apparently paradoxical phenomenon, we combined our electrophysiological data with a well-established biophysical CA1 network model (Fig. 7a) of theta-nested γ-oscillations^31–33^. In brief, the model is composed of Hodgkin-Huxley-type neuron models of O-LM (“O-cells”; inhibitory), pyramidal (“E-cells”; excitatory), and fast spiking PV^+^ cells (“I-cells”; inhibitory). We modeled the E-population as a single cell firing at the population frequency^31, 32^, thereby generating EPSPs in the I-cells (n=10) and O-cells (n=10) at gamma frequency, as observed experimentally^34^.

**Figure 7.**
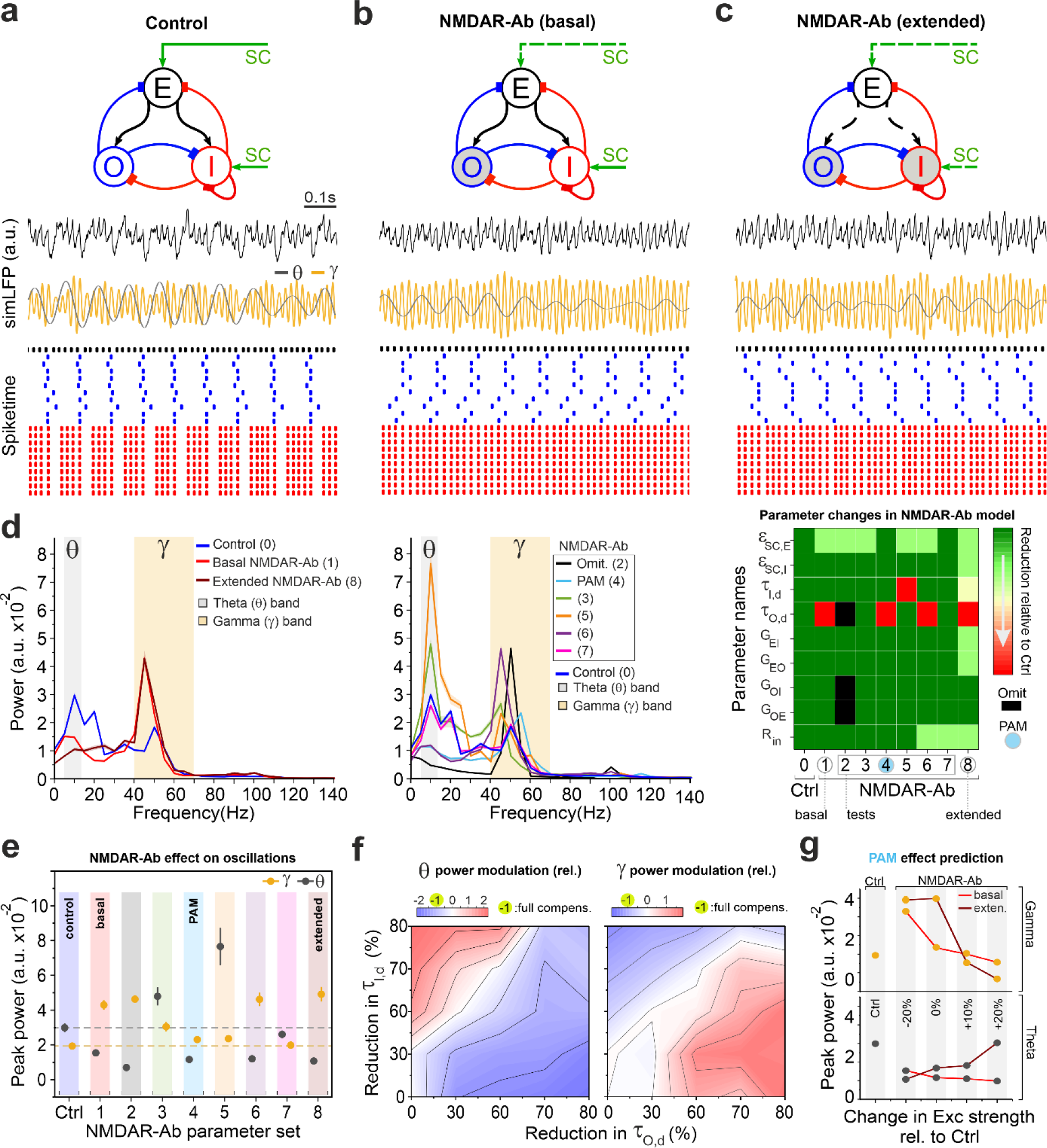
A CA1 network model identifies disinhibition of PN-PV^+^ subnetwork as underpinning gamma amplification under NMDAR-Ab. **(a)** The model reproduces the hippocampal theta-nested gamma oscillations, using Control parameterization. *Top panel*: Schematic diagram representing the synaptic connections among three distinct cell populations. E: pyramidal cells (excitatory), I: fast spiking PV^+^ cells (inhibitory), O: O-LM cells (inhibitory), SC: Schaffer collaterals delivering excitatory input to E-and I-cells. *Second panel*: Example LFP signal simulated by the Control network model (simLFP), shown for one (out of ten) second. *Third panel*: corresponding band-pass filtered simLFPs in theta and gamma frequency bands. *Bottom panel*: Corresponding spike-rastergram of the model underlying the depicted simLFP. **(b)** The dysfunction of O-cells under NMDAR-Ab decreased θ-oscillations and increased γ-oscillations profoundly. Same as (a), but for NMDAR-Ab basal model, which considers the reduction in SC’s excitatory drive to E-cells *ε_SC_*_,*E*_ (coded by dashed line) and in the decay time-constant of synaptic projection of O-cells *τ_O_*_,*d*_ (coded by gray color). **(c)** The main modeling results are robust against extrapolation of NMDAR-Ab effect on synaptic and membrane characteristics. Same as (b), but including a reduction in *ε_SC_*_,*I*_, in *τ_I_*_,*d*_, in Rin of E-cell, and in the synaptic weights of E- to I- and O-populations (*G_EI_* and*G_EO_*). Similar results were obtained as in (b). **(d)** *Left*: Power spectrum of simLFPs shown in (a)-(c). *Middle*: Same as *Left*, but for different parameterizations of the NMDAR-Ab model. *Right*: Look-up map of parameter sets (1-8) of the changes used in NMDAR-Ab model, relative to Control model (0). Exact values can be found in Supplementary Table 2. The mean (solid line) ± SEM (shaded area). **(e)** *Left*: Peak power of θ-oscillations and γ-oscillations for the parameter sets shown in look-up map in (d). **(f)** Compensatory role of faster inhibitory kinetics after NMDAR-Ab treatment. The color-coded contour-maps encoding the amount of change in the increased θ- (left) and γ-oscillatory powers (right) of model #3 in (d), through acceleration of synaptic inhibition. Value of -1 encodes full suppression of the excessive powers. The reductions (i.e. acceleration) of *τ_I_*_,*d*_ and *τ_O_*_,*d*_ are in relative to Control model. **(g)** Augmenting Exc synaptic strength can reinstate γ- (upper panel) and boost θ- oscillatory (lower) powers. The changes in synaptic strengths are in relative to Control model; see also Supplementary figure 5b.

Figure 7a shows a typical simulated LFP signal (simLFP) in the Control model, emulating the reported theta-nested γ-oscillations in CA1^30, 31^. The spike-rastergram shows that O-cells fire preferentially and coherently at theta frequency and modulate the cycles of γ-oscillations produced by the E-I subnetwork. We next investigated the key neural alterations underlying the aberrant γ-oscillations by re-parameterizing the Control model. Our analysis revealed that implementing just two of the major changes seen in NMDAR-Ab group can sufficiently account for the emerged aberrant oscillations (Fig. 7b): the reduction in SC synaptic drive to E-cells, *ε_SC_*_,*E*_, reflecting reduced synaptic AMPAR signaling, and the reduced decay time constant of synaptic inhibitory projection of O-cells, *τ_O_*_,*d*_, reflecting faster decay in GABAAR currents. Under these two changes (NMDAR-Ab basal model), we found that γ-oscillatory power is increased strongly (Fig. 7b,d), while its peak frequency is closely preserved (∼48-49 Hz). Moreover, O-cells’ firing coherence was largely lost at the population level, leading to a decrease in θ-oscillatory power (Fig. 7b; see blue spiketimes). These results corroborate our experimental data (Figs. 6). Importantly, inspection of spike-rastergrams together with simLFPs (Fig. 7b) shows that O-cells effectively fail in nesting the gamma rhythm, thereby causing a disinhibition of E-I subnetwork, which in turn increases the amplitude and temporal continuity of γ-oscillations. We further confirmed this finding by omitting the O-population from the NMDAR-Ab basal model, where θ-oscillations, as expected, were largely abolished, but importantly, γ-oscillations underwent a similar amplification (compare models 1 and 2; Fig. 7d,e). Mechanistically, these results suggest that disinhibition of the γ-oscillations’ generator (E-I subnetwork), caused by O-cells’ dysfunction, underlies the increased γ- oscillations in the NMDAR-Ab group.

By applying the change in either *ε_SC_*_,*E*_ or *τ_O_*_,*d*_ separately to the NMDAR-Ab basal model, we further found that neither of these changes individually, but instead their combination, can reliably reproduce our measured LFPs (models 3 and 4; Fig. 7d,e). Reducing Rin in E-cells had a negligible effect (models 6 and 7 in Fig. 7d,e). Moreover, reducing decay time constant of fast inhibition mediated by I-cells (*τ_I_*_,*d*_), instead of *τ_O_*_,*d*_, in NMDAR-Ab basal model failed to reproduce our measured LFPs (model 5; Fig. 7d,e). This implies that from the overall reduction in τ_*decay*_ of the GABAA synapses onto CA1-PNs under NMDAR-Ab (Fig. 3), including those from PV-cells (Fig. 4), the reduction in τ_*decay*_ from O-LM cells is the sufficient condition to account for the observed aberrant γ-oscillations. In sum, these results suggest that the combined reductions in the time-course of slow synaptic inhibition and the SC’s excitatory drive to E-cells are the necessary and sufficient factors for γ-oscillations amplification in NMDAR-Ab group.

Theoretical studies predicted that faster inhibitory kinetics can counteract the disturbed network stability following NMDAR-hypofunction^35^ . Our modeling reaffirmed this by showing that the excessive θ- and γ-oscillatory activity emerging under NMDAR-Ab-induced reduction in *ε_SC_*_,*E*_ (model 3 in Fig. 7d,e) can be suppressed by reductions in *τ_O_*_,*d*_ and *τ_I_*_,*d*_, respectively (Fig. 7f, and models 1 and 5 in Fig. 7d,e). Corroborating theoretical predictions^35^, these results thus designate a compensatory role for the faster inhibitory kinetics observed after NMDAR-Ab treatment (Figs. 3,4; see also Discussion).

To investigate the significance of other plausible or measured neural alterations under NMDAR-Ab, we also extended our NMDAR-Ab basal model (Fig. 7c): I) We reduced the synaptic strength of all excitatory connections within CA1. II) We reduced not only *τ_O_*_,*d*_ but also *τ_I_*_,*d*_ (Fig. 3). III) We reduced Rin of E-cell (Fig. 2a-d). This extended model again exhibited an increase in γ-oscillations and a suppression of θ-oscillations (Fig. 7c,d,e). These results demonstrate the robustness of our modeling results, and also underscore the modulatory, rather than essential, role of these extensions in mediating γ-oscillation amplification.

Finally, we found that removing the reduction of *ε_SC_*_,*E*_ in NMDAR-Ab basal model can effectively reinstate γ-oscillatory power (model 4; Fig. 7d,e). As also confirmed in the NMDAR-Ab extended model, these results predict that sufficiently augmenting excitatory synaptic strength in the NMDAR-Ab group can markedly suppress the excessive γ-oscillations (Fig. 7g, Supplementary figure 5).

### AMPA-PAM alleviates abnormal γ-oscillations

We next aimed at experimentally testing not only our model predictions on modulation of AMPAR signaling (Fig. 7g) but also whether the γ-oscillations’ amplification (Fig. 6) is present in persistent network oscillations. For this purpose, we first chemically stimulated hippocampal slices with 20µM carbachol (CCH) and recorded LFP signals from CA1 pyramidal layer^36^. CCH induced γ-oscillations in both groups, again showing a profound increase in their power under NMDAR-Ab, while preserving peak frequency at ∼25 Hz (Fig. 8a, b). Moreover, by computing the time at which the envelop-autocorrelation of LFP signals drops below 0.5 value (‘0.5-Lag’; Fig. 8d,e)^37, 38^, we found an aberrant, higher autocorrelation of γ-oscillations over longer time scales under NMDAR-Ab, as quantified by its higher 0.5- Lag values (Fig. 8d). This elevated autocorrelation under basal conditions can be detrimental to e.g. the discrimination and retention of memory traces^39^.

**Figure 8.**
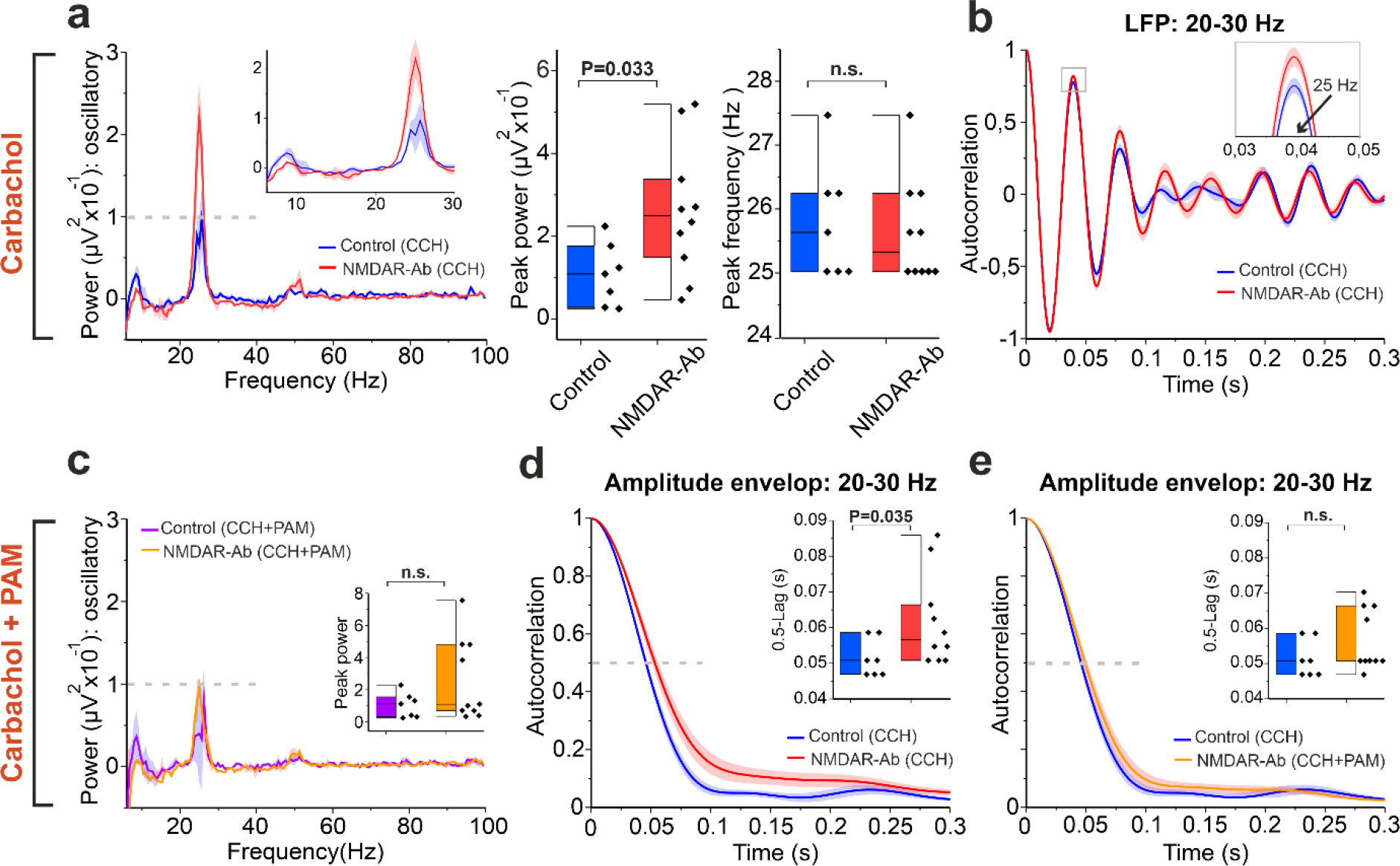
AMPA-PAM suppresses the NMDAR-Ab-induced amplification of persistent γ-oscillations in CA1. **(a)** *Left*: Power spectrum of 10-minutes recordings of LFPs (2-251 Hz) from CA1 hippocampal slices under carbachol (CCH) application. *Inset*: Zoom-in of the power spectrums at lower frequency ranges. Note the abnormal, higher power of oscillations in ∼20-30 Hz under NMDAR-Ab (n=10 slices) as compared to Control (n=7), and a virtually opposite effect on theta band (5-10 Hz). The median (solid line) ± jackknife standard error of median (shaded area). *Middle*: Peak power of γ-oscillations within 20-30 Hz range. *Right*: Same as *Middle*, but for the peak frequency of γ-oscillations. **(b)** Autocorrelograms of the LFPs band-pass filtered between 20-30 Hz. *Inset*: Zoom-in of the gray square. **(c)** AMPA- PAM tends to suppress the abnormal, excessive γ-oscillations power caused by NMDAR-Ab. Same as (a), but for the signals recorded under the application of AMPA-PAM, in addition to CCH. Note AMPA-PAM pushes the power γ-oscillations power under NMDAR-Ab towards that under Control (i.e. to ∼0.1 µV^2^; dashed lines in (a) and (c)). *Inset*: Same as (a, *Middle*), but under CCH+PAM application. **(d)** NMDAR-Ab causes an aberrant prolongation of the temporal correlation of γ-oscillations. Autocorrelograms of the amplitude envelop of LFPs band-pass filtered between 20-30 Hz. The dashed line designates the time that the autocorrelation drops below 0.5 value (0.5-Lag time). *Inset*: Box plot of the 0.5-Lag times. (e) Same as (d), but for Control (CCH) and NMDAR-Ab (CCH+PAM). See Supplementary Table 1 for statistical details.

Finally, we experimentally tested our model prediction about restoring γ-oscillations through augmenting the reduced excitatory synaptic strength (see also Fig. 7g). We repeated the LFP recordings under the application of both CCH and a selective positive allosteric modulator (PAM) of AMPARs (10µM LY404187) to enhance AMPA signaling pharmacologically^40^. We found that PAM, on average, suppresses γ-oscillations’ peak power in NMDAR-Ab group (note the dashed line at ∼0.1µV^2^ in Fig. 8a,c), hence confirming our model prediction. Moreover, PAM was able to restore the normal correlation level in NMDAR-Ab group (Fig. 8e), while not exerting a significant change in Control-Ab group. Therefore, the effective restoration of γ-oscillations’ characteristics by AMPA-PAM provides a novel, target-directed rescue strategy in NMDAR-Ab encephalitis.

## Discussion

NMDARs are critical for brain function being involved in synaptic plasticity^41^, homeostatic scaling^42^, neuronal spike-timing^43^, and persistent network activity^44, 45^. Here, we provide in- depth insights into the open question of how specific NMDAR autoantibodies (Abs) induce widespread, severe cognitive and behavioral deficits. We specifically focused on potential impairment of E-I-balance and network oscillations as important determinants of network and cognitive functions^46, 47^. This follows as these mechanisms were altered in disease states such as schizophrenia^48, 49^, to which NMDAR-encephalitis shares many similarities. Using proteomic, synaptic, cellular, and network-level measurements in the hippocampal CA1, we reveled a dual effect of pathogenic NMDAR-Ab: Whereas Abs induce synaptic and cellular hypoexcitability, they render network activity hypersynchronous in the gamma-band, with disrupted theta-gamma coupling.

We extensively investigated the hippocampal CA3→CA1 feed-forward microcircuit, in which E-I-balance dynamics directly regulate the timing and rate of PN activity^17, 18, 50, 51^. NMDAR-Ab- induced hypoexcitability of CA1-PN, due to reduced synaptic (AMPA) and intrinsic excitability, decreased AP jitter, which in turn may disturb both rate and spike-time coding processes^52^ in the CA1. Furthermore, we found the STP of E-I-balance to be perturbed by NMDAR-Ab, leading to a reduced excitatory integration window which can limit coincidence detection^53^ and dynamic range for excitatory inputs onto CA1-PNs^51, 54^. This induced STP perturbation may also account for the reduced serial dependence in NMDAR-encephalitis and schizophrenia patients^22^.

We also found a plethora of dysregulated proteins in the hippocampus after NMDAR-Ab treatment. Virtually all major components of glutamatergic signaling were affected (AMPAR, NMDAR, KAR, metabotropic receptor groups I, II and III), as well as central hubs of intracellular signaling (including CaN, CaMK, PKA, PKC, Stat1), raising the prospect of metaplasticity alterations. How NMDAR-Ab induce these changes, and whether they are primary or compensatory, remains to be determined. However, the changes in AMPAR and KAR were also observed after chronic phencyclidine treatment^55^, indicating that NMDAR play a primary role in resetting the strength of glutamatergic signaling. The specific effect of NMDAR-Ab on LTP pathways suggests that the changes in AMPAR (and AMPAR- associated proteins) result from a direct impairment of NMDAR-dependent homeostatic mechanisms. Strikingly, we also found several proteins strongly dysregulated in schizophrenia to also be modified by NMDAR-Ab, highlighting possible common disease mechanisms (Cacna1c, Dlg1, Esyt1, Synpo, Dnm3, Shank^56^, etc.).

Identifying the specific neural alterations underpinning the abnormal gamma amplification is important for understanding network dysfunction and developing treatment strategies. γ- oscillations typically arise from the interaction between PNs and INs, whose synaptic current kinetics can modulate the power and frequency of these oscillations^27, 29, 57^. Accordingly, our findings about reduced AMPAR-mediated drive and faster kinetics of GABAAR signaling point to effective alteration of γ-oscillations characteristics. Furthermore, our observed dominance of inhibition and the reduced EPSP and AP jitter can also promote the network hypersynchronicity^58^. By integrating our synaptic and cellular data into a biophysical CA1 network model, we found that neither of these changes individually, but instead their combination, can reliably emulate the oscillation characteristics in NMDAR-Ab group. Of note, our recorded aberrant gamma activity is in agreement with multiple studies based on genetic or pharmacologic manipulation of NMDAR, showing a robust alteration of γ- oscillations due to NMDAR-hypofunction (for a review see^2^).

Using the model, we further found that these NMDAR-Ab-induced biophysical changes should mainly impair INs projecting slow, rather than fast, GABAergic synapses (here, OLM and PV cells, respectively). This in turn brings about the disinhibition of CA1’s PN-PV gamma generation subnetwork (the so-called PING network^32^), hence increasing γ-oscillatory power. Our modelling results imply that this disinhibition is mainly due to the failure of OLM-cells in providing effective inhibition onto the PING network. This finding is supported by experimental evidence showing that inhibition arising from these somatostatin positive interneurons (SOM-IN) onto PNs (and PVs) has a pivotal role in confining the excitatory net effect of Schaffer collateral (CA3→CA1), temporoammonic (EC→CA1), and hippocampal output pathways^59–61^. Remarkably, our model predicted that augmenting AMPAR signaling can restore the normal γ-oscillations. Indeed, we showed that bath-application of AMPA-PAM is able to effectively suppress the excessive gamma in the NMDAR-Ab group, thereby confirming this prediction.

OLM cells also play a key role in the generation of θ-oscillations in hippocampus^34, 62^. Our measurements and network modelling provide evidence for the alteration of these oscillations under NMDAR-Ab. Impaired θ-oscillations were also found after NMDAR-ablation in PV-IN^11^, as well as due to disruption of OLM cells in an epilepsy model^33^, or through impaired inhibition onto PV-IN^31^.

The purpose of these synaptic changes resulting in aberrant γ-oscillations is an open question. This may be addressed by considering the experimentally-supported role of relatively slow NMDAR-mediated currents in stabilizing global network activity and in the emergence of stable (asynchronous) persistent working memory states^35, 44, 45, 63, 64^. Conceptually, NMDAR reduction can cause fast excitation to outpace reverberant inhibition, hence making it more prone to instability (e.g. seizure-like activity), hypersynchronous (here, in network oscillations), or self-driven oscillatory dynamics^35, 65^. Similarly, an impaired NMDAR-driven stability may explain data from CA3 after NMDAR-Ab treatment or region- specific deletion of NMDARs^66, 67^. The induced aberrant oscillations can be detrimental to network stability behavior and function^65, 68^.

Theoretical studies predicted two mechanisms amenable to compensate for the disturbed NMDAR-driven network stability, namely weakening of AMPAR strength and acceleration of GABAAR synaptic inhibition^35, 45^. Whereas these predictions were mainly derived based on cortical neural networks, our findings suggests a pathological role of reduced AMPAR strength in CA1. This is because our CA1 network model and experimental data found AMPA-PAM to normalize hippocampal rhythmogenesis in NMDAR-Ab group. In contrary, these data implies a homeostatic role of faster inhibition in a rhythm- and interneuron-type- selective manner: The NMDAR-Ab-affected CA1 network may prioritize alleviating the aberrant θ-oscillations over γ-oscillations by triggering a stronger homeostatic reduction in *τ_O_*_,*d*_ relative to *τ_I_*_,*d*_ . Whereas this effectively suppresses excessive θ-oscillations, it concurrently renders γ-oscillations disinhibited. Nonetheless, the applicability of these findings, including the therapeutical effect of AMPA-PAM in cortical networks, requires future investigations.

Interregional communication as occurs e.g. during execution of working memory or learning has been linked to gamma-oscillations, which are seen as a synchronization mechanism enabling functional connectivity^53, 69^. Theta s mechanistic insights into the pathophysiology of neuropsychiatric symptoms in NMDAR-encephalitis and propose novel therapeutical perspectives. We conclude that NMDAR-Abs disrupt the E-I-balance and induce cellular hypoexcitability which, together with faster inhibitory kinetics, causes aberrant oscillatory dynamics in the hippocampus.

## Acknowledgments

We thank Albert Compte for stimulating discussions and very constructive comments on this manuscript. We thank Claudia Sommer, Christin Reißig, Marc Raugust, and Marin Kempfer for expert technical assistance and for image acquisition and analysis. We would like to thank Emil Ceanga for support and very helpful methodological discussions. The authors gratefully acknowledge support from the FLI Core Facility Proteomics. The FLI is a member of the Leibniz Association and is financially supported by the Federal Government of Germany and the State of Thuringia. This work was supported by the German Research Foundation (FOR3004; GE2519/8-1, GE2519/9-1, to C.G.; Research Training Group ProMoAge GRK 2155 to A.O.), by the German Federal Ministry of Education and Research (01GM1908B, 01EW1901 to C.G.), by the Interdisziplinäres Zentrum für Klinische Forschung (IZKF) Jena (to M.C, H.H.), the Foundation „Else Kröner-Fresenius-Stiftung“ within the Else Kröner Research School for Physicians „AntiAge“ (to M.C.) and award number: 2019_A79 (to A.O.), the Fritz-Thyssen foundation (award number: 10.20.1.022MN to A.O.), the Chan Zuckerberg Initiative Neurodegeneration Challenge Network (award numbers: 2020-221617 and 2021-230967 to A.O.) and the Schilling Foundation (to C.G.). Schematic figures of experiment set-ups have been created using Biorender.com.

## Author contributions

M.C., V.R., and C.G conceptualized the study; M.C. and H.H. conducted electrophysiological recordings; M.C. and V.R. performed electrophysiological data analysis; V.R. conducted the implementation and analysis of network modeling; A.O. conducted and analyzed proteomics data; L.S. performed morphological analysis; J.K. and H.P. conducted antibody synthesis and purification experiments; M.C., V.R., C.G., D.H., J.K., H.P., L.G., S.H., J.D., A.O. and M.H. performed data interpretation; M.C., V.R. and C.G. wrote the manuscript with inputs from all authors; C.G. supervised this study.

## Data and code availability

Statistical data associated with this study is presented in Supplementary Table 1. All datasets generated during this study are available from the corresponding authors upon reasonable request. Mass spectrometry proteomics data have been deposited to the ProteomeXchange Consortium via the PRIDE^70^ partner repository.

Code associated with the neural network model is publicly available code in ModelDB (https://senselab.med.yale.edu/ModelDB; accession number: 138421). The Matlab code for IRASA method can be found at https://purr.purdue.edu/publications/1987/1.

## Online Methods

See the attached document.

## Online Methods

### Animal model of NMDAR-encephalitis

Forty male C57BL/6J mice were housed in a room maintained at a controlled temperature of (21±1°C) and humidity (55±10%) with illumination at 12-hour cycles; food and water were available *ad libitum*. Animal experiments were performed in accordance with the ARRIVE guidelines for reporting animal research^1^ and the experimental protocol was in accordance with European regulations (Directive 2010/63/EU) and was approved by the local ethics committee at the University of Jena. Sixteen-weeks old (25-30g) mice were implanted with biventricular osmotic pumps (model 1002, Alzet, Cupertino, CA) with the following characteristics: volume 100 µl, flow rate 0.25 µl/hour, and duration 14 days, as previously reported^2, 3^. The day before surgery the two pumps were each filled with 100µl of 1µg/µl human monoclonal NMDAR-Ab (#003-102) or Control-Ab (#mGO53)^4^. Mice under isoflurane anesthesia were placed in a stereotactic frame, and a bilateral cannula (model 3280PD2.0/SP, PlasticsOne) was inserted into the ventricles (coordinates: 0.2 mm posterior and ±1.00 mm lateral from bregma, depth 2.2 mm). The cannulas were connected to two subcutaneously implanted osmotic pumps on the back of the mice.

### Electrophysiological Recordings

#### Acute hippocampal preparation

Two weeks (14-16 days) after intraventricular osmotic pump implantation, mice were deeply anesthetized with Isoflurane, decapitated, and the brain was removed in ice-cold protective artificial cerebrospinal fluid solution (paCSF) containing, in mM: 95 N-Methyl-D-Glucamine, 30 NaHCO3, 2.5 KCl, 1.25 NaH2PO4, 10 MgSO4, 0.5 CaCl2, 20 HEPES, 25 glucose, 2 thiourea, 5 Na-ascorbate, 3 Na-pyruvate, 12 N-acetylcysteine, adjusted to pH 7.3 and 300- 310 mOsmol, saturated with Carbogen. Hippocampal 350µm-thick transverse slices were prepared on a vibratome (VT 1200S, Leica, Wetzlar, Germany) in ice-cold paCSF and transferred for warm recovery in paCSF at 32°C for 12 minutes. Before recordings, slices were left to recover for at least 60 minutes at room temperature in aCSF+ containing in mM: 125 NaCl, 25 NaHCO3, 2.5 KCl, 1.25 NaH2PO4, 1 MgCl2, 2 CaCl2, 25 glucose, 2 thiourea, 5 Na-ascorbate, 3 Na-pyruvate, 12 N-acetylcysteine, adjusted to pH 7.3 and an osmolarity of 300–310 mOsmol, and saturated with Carbogen. For recordings, slices were transferred in a recording chamber under continuous perfusion (2-4 ml/min, Ismatec, Wertheim, Germany) with 27°C aCSF containing in mM 125 NaCl, 2.5 KCl, 25 NaHCO3, 1.25 NaH2PO4, 1 MgCl2, 2CaCl2, saturated with Carbogen.

#### Whole-cell recordings

Researchers performing all recording experiments and analyses were blinded to the experimental treatment of the animals. CA1 pyramidal neurons (CA1-PN) were visually identified using a microscope (Examine.Z1, Zeiss, Jena, Germany) equipped with differential interference contrast optics. Patch pipettes were pulled using a P-87 horizontal pipette puller (Sutter Instruments, Novato, CA, USA) from thick-walled borosilicate glass (0.86x1.50, Science Products, Kamenz, Germany) and had a resistance of 2.5-5 MΩ when filled with intracellular solutions. For voltage-clamp recordings, the intracellular solution contained in mM 135 CsMeSO4, 3 CsCl, 10 HEPES, 0.2 EGTA, 0.1 spermine, 2 QX-314-Br, 2 Mg2-ATP, 0.3 Na2-GTP, 10 phosphocreatine (pH 7.25, osmolarity 280 mOsmol), while for current-clamp recordings the intracellular solution contained in mM 150 K-Gluconate, 1.5 MgCl2, 10 HEPES, 0.2 EGTA, 0.1 spermine, 2 Mg2-ATP, 0.3 Na2-GTP, 10 phosphocreatine (pH 7.25, osmolarity 280 mOsmol). For morphological reconstructions 3mg/ml biocytin was freshly added to the intracellular solutions before recordings. During voltage-clamp recordings, 1µM AM-251 was added to aCSF to block depolarization-induced suppression of inhibition ^5^. For field potential measurements, the recording electrode was filled with aCSF.

Voltage-clamp (Vc) and current-clamp (Ic) signals were recorded and digitized with an EPC 10 USB double amplifier/digitizer (HEKA, Ludwigshafen/Rhein, HRB 41752) or recorded with a MultiClamp 700B amplifier (Molecular Devices, Sunnyvale, CA, USA), and digitized using an Axon Digidata 1550B digitizer (Molecular Devices, Sunnyvale, CA, USA). Signals were low-pass filtered at 2kHz (Vc) or 5kHz (Ic) and digitized at 20kHz. Data acquisition was performed in PatchMaster v2x80 (HEKA, Ludwigshafen/Rhein, HRB 41752) or Clampex 10.7 (Molecular Devices, Sunnyvale, CA, USA). Measured and applied potentials were corrected online for an experimentally determined liquid junction potential of 8mV for both intracellular solutions. Recording of the resting membrane potential was performed immediately after break-in, and cells with a membrane potential higher than -60mV were discarded. Series resistances (<25MΩ) was 70% compensated and was monitored throughout recordings. Cells were discarded if a >20% change in series resistance occurred. Analysis of recorded signals was performed in Clampfit 10.7 (Molecular Devices, Sunnyvale, CA, USA) or in Matlab (MathWorks) using custom-written code (see below).

Stimulation of the Schaffer collaterals (SC) was performed with an aCSF-filled micropipette acting as a monopolar stimulation electrode that was placed 200–300µm away from the recording electrode. Stimulation was applied through a constant current stimulation unit (DS3, Digitimer) for 40µs at an intersweep frequency of 0.1Hz. The reversal potentials of synaptic excitation and inhibition were measured after bath application of 100µM PTX (for *Eexcitation*) or 10µM CNQX and 50mM AP-5 (for *Einhibition*), after which CA1-PN were recorded at different holding potentials during single SC stimulation (Supplemental Fig. 2a). The reversal potentials were measured as -5 mV and -70 mV for excitation and inhibition, respectively.

To determine the synaptic input/output (I/O) curves of CA1-PN, recorded cells in Ic mode were held at 0pA while SC were stimulated at incremental strengths at 0.1Hz until action potential (AP) probability reached 100% (ca. 80-100 sweeps). EPSP slopes were binned, and AP probability was determined per bin^6^. To determine the input resistance, we calculated the slope of a straight-line fit to the voltage-current curve which was established using subthreshold current injections. The membrane time constant was measured by fitting a single exponential to the voltage response of a -20pA current step. Voltage sag and APs were elicited by current step injections from -240pA to 360pA in 20 pA steps. AP threshold was defined as the potential at which the rise slope exceeded 15V/s^7^. The spike frequency adaptation was calculated as the ratio of the last interspike interval to the first one.

The impedance amplitude profile (ZAP) was determined by injecting a 20s-long, linearly increasing chirp current (0-15Hz) from a baseline of 0pA (at resting potential). The amplitude of the injected current was adjusted to evoke a subthreshold, ∼10mV peak-to-peak response. Following previous studies^8, 9^, we calculated ZAP as a function of the input frequency *Z* (*f*) = V^-^ (f) *I*^-^ (*f*), where V^-^ (f) and *I*^-^ (*f*) are the fast Fourier transform (fft) of the cell membrane potential *V* and the applied chirp current *I*. Both *V* and *I* were downsampled to 2 kHz prior to fft. *Z* (*f*) is a complex number and its absolute value determines the impedance amplitude at frequency *f*, denoted as *Z* (*f*) . In our results we refer to *Z* (*f*) as the impedance. For each cell, we computed averaged across the trials. *Z* (*f*) for each sweep (out of 3) separately, then To evoke biphasic (eEPSC-eIPSC) and monophasic (eEPSC or eIPSC) responses, SC were incrementally stimulated and eEPSCs were recorded at a holding potential of -70mV until a plateau potential was reached (eEPSC’s I/O curve). Thereafter, supramaximal stimulation intensity was used. Biphasic responses were elicited by changing the holding potential to approx. -35mV and optimizing the holding potential for a maximal amplitude of both components (range -40mV to -25mV). Short-term plasticity was investigated by applying paired-pulse supramaximal stimulation of the SC at 50ms and 200ms intervals, and train stimulation by applying a 100-pulse train of supramaximal stimulation of the SC at 20Hz. The excitatory integration window was measured as the time interval between the peak of the excitatory (eEPSC) and the peak of the inhibitory (eIPSC) components in biphasic responses.

Local field potentials (LFP) were recorded only using a MultiClamp700B with the headstage in voltage-follower mode, with a low-pass Bessel filter of 1kHz and a sampling frequency of 20kHz. For electrically evoked oscillations, the recording electrode, consisting of a micropipette (∼2MΩ) filled with aCSF, was placed in the stratum pyramidale of the hippocampal CA1 region, while the stimulus electrode was placed 200-300µm away in the stratum radiatum towards the CA3. Supramaximal stimulation (1000µA) of the SC at 100Hz was repeated for three sweeps (60s intersweep interval). For chemically evoked oscillations, baseline LFP was measured for 5 minutes, before adding 20µM carbachol or 20µM carbachol with 10µM LY404187 (AMPA-PAM) to the bath solution, after which LFP was measured for 7 minutes.

Analysis of synaptic currents was performed in Clampfit 10.5 using the template search function. Templates of spontaneous excitatory and inhibitory postsynaptic currents (sEPSC and sIPSC, respectively) were obtained by averaging 25-35 manually-selected events per group. All events with a detection similarity threshold of three were further verified by visual inspection. Decay time constants were estimated by a single exponential fit. The median was used to calculate the central tendency of synaptic parameters per cell (sEPSC/sIPSC amplitude, frequency, decay tau, charge transfer).

#### Biocytin staining and morphologic analysis

After recording, slices were transferred in 4% PFA and fixed overnight at 4°C. After washing in TBS, slices were incubated for two hours in TBS plus (TBS, 5% donkey serum, 0.3% Triton-X-100) and then incubated overnight in 1:500 Streptavidin-Cy3 (Sigma) in TBS plus before embedding in Entellan (Merck, Darmstadt, Germany). Filled neurons were captured on a confocal scanning microscope (LSM 710, Zeiss, Jena, Germany) and morphologic analysis was performed in Imaris7 (Bitplane, Belfast, UK).

### Sample preparation for proteomics

Immediately after decapitation, several hippocampi were dissected and flash-frozen at -40°C and subsequently stored at -80°C. Hippocampi were thawed and transferred into Precellys® lysing kit tubes (Keramik-kit 1.4/2.8 mm, 2 ml (CKM)) containing 200 µl of PBS supplemented with 1 tab of cOmplete™, Mini, EDTA-free Protease Inhibitor per 50 ml. For homogenization, tissues were shaken twice at 6000 rpm for 30 s using Precellys® 24 Dual (Bertin Instruments, Montigny-le-Bretonneux, France) and the homogenate was transferred to new 1.5 ml Eppendorf tubes. For each sample, 100 µl of homogenate was diluted with 2x lysis buffer (final concentrations: 2 % SDS, 100 mM HEPES, pH 8.0), sonicated in a Bioruptor Plus (Diagenode, Seraing, Belgium) for 10 cycles with 1 min ON and 30 s OFF with high intensity at 20 °C, and then heated at 95 °C for 10 min. A second sonication cycle was performed as described above. Lysates were stored at -80 °C until further processing.

Lysates were thawed, sonicated as described above, and aliquots corresponding to 100 µg of protein were reduced using 10 mM DTT for 30 min at room temperature and alkylated using freshly made 15 mM IAA for 30 min at room temperature in the dark. Subsequently, proteins were acetone precipitated and digested using LysC (Wako sequencing grade) and trypsin (Promega sequencing grade), as described by Buczak *et al.*^10^. The digested proteins were then acidified with 10 % (v/v) trifluoracetic acid and desalted using *Waters Oasis® HLB µElution Plate 30 µm* following manufacturer instructions. The eluates were dried down using a vacuum concentrator and reconstituted in 5 % (v/v) acetonitrile, 0.1 % (v/v) formic acid. For mass spectrometry (MS) analysis, samples were transferred to an MS vial, diluted to a concentration of 1 µg/µl, and spiked with iRT kit peptides (Biognosys, Zurich, Switzerland) prior to analysis.

### Proteomics data acquisition

Approximatively 1μg of reconstituted were separated using a nanoAcquity UPLC (Waters, Milford, MA) was coupled online to the MS. Peptide mixtures were separated in trap/elute mode, using a trapping (Waters nanoEase M/Z Symmetry C18, 5μm, 180 μm x 20 mm) and an analytical column (Waters nanoEase M/Z Peptide C18, 1.7μm, 75μm x 250mm). The outlet of the analytical column was coupled directly to an Orbitrap Fusion Lumos mass spectrometers (Thermo Fisher Scientific, San Jose, CA) using the Proxeon nanospray source. Solvent A was water, 0.1% formic acid and solvent B was acetonitrile, 0.1% formic acid. The samples were loaded with a constant flow of solvent A, at 5 μL/min onto the trapping column. Trapping time was 6 min. Peptides were eluted via the analytical column with a constant flow of 300 nL/min. During the elution step, the percentage of solvent B increased in a nonlinear fashion from 0% to 40% in 120 min. Total runtime was 145 min, including cleanup and column re-equilibration. The peptides were introduced into the mass spectrometer via a Pico-Tip Emitter 360 µm OD x 20 µm ID; 10 µm tip (New Objective) and a spray voltage of 2.2 kV was applied. The capillary temperature was set at 300 °C. The RF lens was set to 30%. Full scan MS spectra with mass range 350-1650 m/z were acquired in profile mode in the Orbitrap with resolution of 120,000 FWHM. The filling time was set at maximum of 20 ms with an AGC target of 5 x 10^5^ ions. Data Independent Acquisition (DIA) scans were acquired with 40 mass window segments of differing widths across the MS1 mass range. The HCD collision energy was set to 30%. MS/MS scan resolution in the Orbitrap was set to 30,000 FWHM with a fixed first mass of 200m/z after accumulation of 1x 10^6^ ions or after filling time of 70ms (whichever occurred first). Data were acquired in profile mode. For data acquisition and processing Tune version 2.1 and Xcalibur 4.1 were employed.

### Proteomics data analysis

Raw DIA files were analysed in directDIA mode using Spectronaut Pulsar (v15, Biognosys, Zurich, Switzerland). The data were searched against a species-specific protein database (Uniprot *Mus musculus* release 2016_01) with a list of common contaminants. The data were searched with the following modifications: carbamidomethyl (C) as fixed modification, and oxidation (M) and acetyl (protein N-term) as variable modifications. A maximum of 2 missed cleavages was allowed. The search was set to 1 % false discovery rate (FDR) at both protein and peptide levels. DIA data were then uploaded and searched against this spectral library using Spectronaut Professional (v.15) and default settings. Relative quantification was performed in Spectronaut for each pairwise comparison using the replicate samples from each condition using default settings, except: Major Group Quantity = median peptide quantity; Major Group Top N = OFF; Minor Group Quantity = median precursor quantity; Minor Group Top N = OFF; Data Filtering = Q value; Normalization Strategy = Local normalization; Row Selection = Automatic; Exclude Single Hit Proteins = TRUE. Differential abundance testing was performed using an un-paired t-test between replicates. P values were corrected for multiple testing multiple testing correction with the method described by Storey^11^. The data (candidate table) and protein quantity data report were exported, and further data analyses and visualization were performed with R (v.4.0.5) and R studio server (v. 2022.02.0) using in-house pipelines and scripts. KEGG pathway and Gene Ontology over-representation analyses were performed with WebGestalt^12^ using the significant protein groups (q<0.05) against a background of all the quantified protein groups. Significantly enriched pathways and GO terms were defined using a cut-off of FDR < 0.05.

### Analysis of electrically induced oscillations

The HFS-induced network oscillations were recorded as described above. The signals were then processed using custom written code in Matlab (MathWorks). For a proper time-frequency analysis, while reducing the potential stimulus-induced edge-like effects (see below), we subjected each HFS signal to the following correction steps and verified the results visually. (1) The baseline of the before-stimulus section of the signal (∼2.15 seconds) was corrected using the robust polynomial fitting method of maximum 7^th^-order. (2) The sharply increasing baseline of the signal, observed right after the stimulus offset (see Results), was removed using a polynomial of 17^th^ order. To avoid the disturbance of the signal dynamics by this high-order polynomial, we applied this fitting to only 2.1 seconds of the after-stimulus signal. Note that we also considered the 10 milliseconds after the last impulse of HFS as a part of stimulus duration, in order to avoid the introduction of stimulus artifact in our results. (3) Using the same procedure as in step (1), was removed the baseline of the remaining (tail) of the signal after stimulus. (4) The 200-milliseconds stimulus artifact was replaced with the mirrored version (along y-axis) of the first 200-milliseconds of the baseline-corrected HFS signal in (2); note, the time is considered as x-axis. (5) Finally, the baseline-corrected HFS signal was obtained by concatenating the resulted signals in (1)-(4). To extract the local field potential (LFP) activity this signal was then digitally band-pass filtered in the range of 1.1-250 Hz using a Kaiser window finite impulse response filter (KW- FIR) with a sharp transition band bandwidth of 1 Hz, stopband attenuation of 60 dB, and passband ripple of 1%. To avoid the potential edge effects of the signal due to the filtering a mirrored version of the whole signal was appended to the beginning and the end of the signal, prior to the filtering. These copies were cut out of the filtered signal afterwards.

To characterize the transient gamma oscillations induced by the HFS we used the time-frequency analysis of the resultant baseline-corrected LFP signals. We performed this analysis separately for the first ∼2.15 seconds part of the signal before stimulus onset (bLFP) and the last ∼7.6 seconds of the signal after the stimulus offset (aLFP). To this end, we convolved each signal with the complex Morlet wavelets of frequencies from 4 to 110 Hz with the increment of 0.5 Hz. As the wavelet scales we used the number of cycles from 5 to 10 (logarithmically-spaced over these frequency levels) thereby effectively accounting for the time-frequency resolution trade-off of the wavelet transforms. For each level of wavelet scale (or frequency), the LFP power at each time point was computed as the squared magnitude of the corresponding coefficient of the wavelet transform at each time-frequency pair; *P_bLFP_* (*t*,*f*) and *P_aLFP_* (*t*,*f*) . The edge effects in the transform results were considered as missing values, and thus discarded. In the case of aLFP signal, we considered the stimulus duration, which we replaced using the mirrored aLFP signal (see above), as a part of aLFP. This enabled us to shift the left edge effect to the onset of stimulus, and thus have the wavelet transform of aLFP almost free of this edge effect. The mirrored part (i.e. of stimulus duration) was then cut out of the wavelet results. To minimize the effect of potential 1/f power scaling phenomenon as well as e.g. the slice- and electrode-specific idiosyncratic characteristics we computed the relative power values:

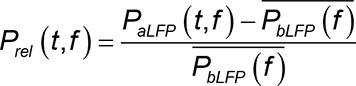

Where, *P_rel_*(*t*,*f*) denotes the induced power after the stimulus at time *t* and frequency level *f*, and *P_bLFP_* (*f*) denotes the time-averaged power of the bLFP at *f*. To investigate the evolution of induced gamma power over time we computed the total instantaneous power as the mean of *P_rel_*(*t*,*f*) over 30-90 Hz (gamma band) at each *t*. To show that our findings are not affected by the normalization we also computed the raw (i.e. non-normalized) total instantaneous power as the mean of *P_aLFP_*(*t*,*f*) over 30-90 Hz at each *t*. To investigate the frequency components of the induced power (i.e. power spectrum) we computed the mean of *P_rel_*(*t*,*f*) over 400 milliseconds after the stimulus offset at each *f*.

Note that this selected window size effectively captures the induced transient gamma responses, before returning to the baseline (see Results). We repeated all these analyses for each sweep (out of three) of the recording session, and then averaged the results over the sweeps to increase the signal-to-noise ratio. For testing the mean difference in gamma power between the two groups (Control vs. NMDAR-Ab) over time, we considered 1.6 seconds of the total instantaneous gamma power time-series after the stimulus offset and divided it to non-overlapping 200-milliseconds bins. We then performed a two-tailed permutation test with 1,000,000 times shuffling at a significance level of 5% while accounting for the multiple-comparison problem by the method of Cohen^13^.

In our preliminary analysis we found that HFS not only induces strong gamma oscillations but also some relatively weak theta responses. Using Chronux software package (http://chronux.org/)^14^, we applied the multi-taper method (time-bandwidth product of 2 with 3 tapers) to the first 500-milliseconds of each aLFP (1.1-250 Hz), in order to first compute the power spectrum of the theta responses. We then assessed how these two responses were coordinated in time using the theta-gamma co-modulation index (TGC)^15^. To this end, we band-pass filtered the baseline-corrected HFS signal using the same KW-FIR filter as above but in the range of 30-90 Hz (for gamma band) and 5-12 Hz (for theta band), separately. This was followed by applying the Hilbert transform and computing the magnitude (i.e. amplitude envelop) of these band-specific signals at each time point, using the resulted analytic signals. This was followed by binning the first 450-milliseconds of these signals using a sliding window of 200 ms size with a step size of 50 ms. The selected bin size was sufficiently large to enclose ∼1-2.5 and ∼6-18 cycles of theta (5-12 Hz) and gamma oscillations (30-90 Hz), respectively (see Results). TGC was then computed as the Pearson correlation between the magnitude time-series of the theta- and gamma-band signals at each bin. To compare the TGC of the groups at the center of each of 5 bins we used the same permutation-test of Cohen^13^ mentioned above.

As an alternative analysis to assess the temporal characteristics of the induced oscillations, we used autocorrelation function (ACF); a measure of the similarity between the values of the same signal over successive time intervals. To do this, for each baseline-corrected LFP signal (5-120 Hz), we computed ACF for 800-milliseconds of its aLFP and of its corresponding bLFP signal preceding the stimulus onset. To compute the induced ACF we then subtracted ACF of the baseline from that of after-HFS.

### Analysis of chemically induced oscillations

The LFP signals under chemical stimulation were recorded as described above. These signals were then processed using custom written code in Matlab (MathWorks). The artifacts were identified by visual inspection and excluded. To extract LFP activity both properly and in a computationally efficient manner, each signal was subjected to the following processing steps. (1) The signal was downsampled to 10 kHz. (2) It was then digitally low-pass filtered using a KW-FIR filter with a passband cutoff of 1 kHz, stopband band of 50 Hz, stopband attenuation of 60 dB, and passband ripple of 1%. (3) This was followed by a downsampling to 5 KHz. (4) Finally, the LFP signal was obtained by band-pass filtering the signal using the same KW-FIR filter as used for HFS data (see above) but in the range of 2-250 Hz.

Field potential signal are usually composed of both fractal (aperiodic or 1/*f* noise) and oscillatory (periodic) dynamics, governed presumably by distinct biological mechanisms. In this work, we are interested in investigating the potential changes in the neural oscillations due to NMDAR-Ab. Hence, in order to separate these components in the power spectrum of the LFP signals we used the Irregular-Resampling Auto-Spectral Analysis (IRASA) method^16^. This enabled us to assess the power characteristics of the oscillatory dynamics in LFP signals exclusively, i.e. without being affected by that of the 1/*f* noise. When applying IRASA to each LFP signal we used a sliding window of 1-second size with 50% overlap. We used the same steps to obtain the LFP signals and the corresponding power spectrums for 10- minutes recorded signals under Carbachol+PAM application.

### Computational network model

To gain mechanistic insights into the mechanism underlying the abnormal increase in gamma oscillations due to NMDAR-Ab we used computational modeling. To do this, we employed a well-established biophysical recurrent neural network model of CA1^17–19^. The model is amenable to reproduce the hippocampal theta-nested gamma oscillations, and has been used extensively in previous studies to explain, or predict, the dynamics of the experimentally observed oscillatory dynamics in LFP signals under various behavioral or pharmacological conditions^17–21^. As compared to the mean-filed CA1 network models^22–24^, this model allows for investigating the network dynamics using a more detailed parameterization of biophysical parameters at both synaptic and single-cell levels. To set the parameter values of this model, we mainly followed previous studies^17, 18^, and considered the resulted model as representing the Control group in our data. We then emulated the NMDAR-Ab condition by applying the relative changes in synaptic and cellular parameter values of the model, according to our measurements (Figs. 1-4). The list of the percentage changes used in the NMDAR-Ab model, relative to Control model, can be found in Supplementary Table 2. By analyzing the simulated LFP (simLFP) and spike rastergrams, we then sought to mechanistically address the mechanism and parameter changes mediating the increase in gamma in NMDAR-AB group.

The model is consisted of the single compartment, Hodgkin-Huxley-type neuron models of pyramidal cells (“E cells”; excitatory), fast spiking PV+ cells (“I cells”; inhibitory), and oriens lacunosum-moleculare interneurons cells (O-LM or “O cells”; inhibitory). Similarly to previous studies^17, 18^, we modeled E-population as a single cell which fires at the population frequency thereby generating EPSPs in the I (n=10) and O (n=10) cells at the gamma frequency, as observed experimentally^25^. In our model, we assumed that the E and I cells receive also external constant (with additive noise) excitatory drive through Schaffer collaterals (SC). However, in general, this drive current can be considered to be delivered by pathways other than SC. In the following, we describe the corresponding neuron and synaptic models, LFP model and its analysis, and the steps used for setting the numeric and random aspects of the model.

### Neuron models

*E cell*: For the pyramidal cell, we use the Olufsen et al. 2003 model^26^ (a variation of Ermentrout and Kopell 1998 model^27^):

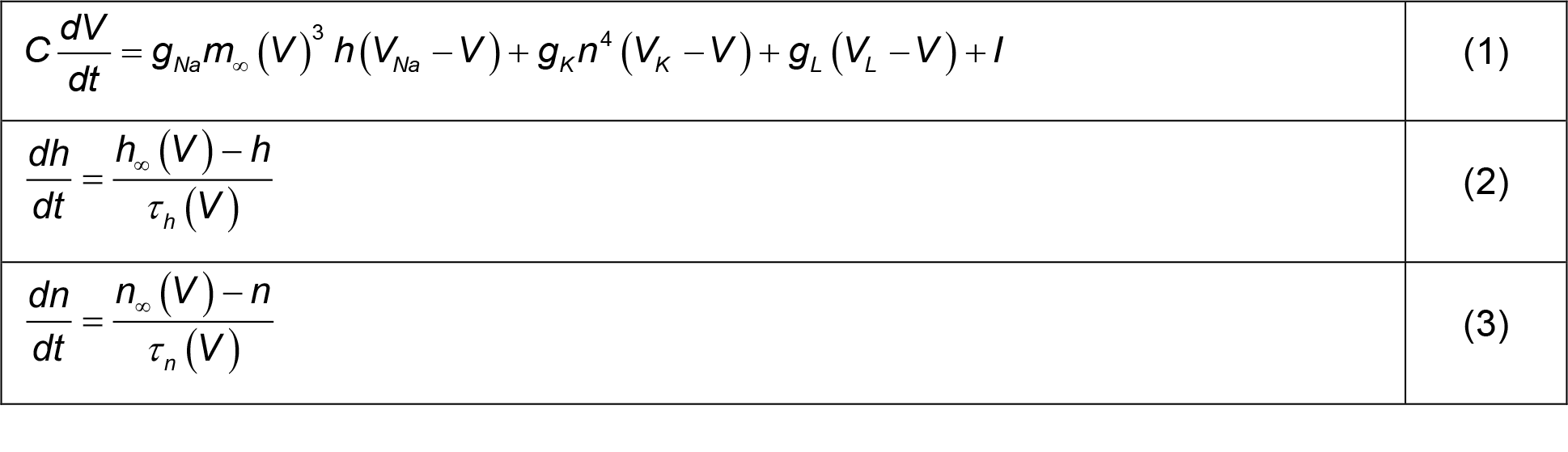

|  |  |
| --- | --- |
| $C \frac{dV}{dt} = g_{Na} m_{\infty}(V)^3 h(V_{Na} - V) + g_K n^4 (V_K - V) + g_L (V_L - V) + I$ | (1) |
| $\frac{dh}{dt} = \frac{h_{\infty}(V) - h}{\tau_h(V)}$ | (2) |
| $\frac{dn}{dt} = \frac{n_{\infty}(V) - n}{\tau_n(V)}$ | (3) |

With

|  |  |
| --- | --- |
| $x_{\infty}(V) = \frac{\alpha_x(V)}{\alpha_x(V) + \beta_x(V)}$ for $x = m, h$, or $n$ | (4) |
| $\tau_x(V) = \frac{1}{\alpha_x(V) + \beta_x(V)}$ for $x = h$ or $n$ | (5) |
| $\alpha_m(V) = \frac{0.32(V + 54)}{1 - \exp(-(V + 54)/4)}$ | |
| $\beta_m(V) = \frac{0.28(V + 27)}{\exp((V + 27)/5) - 1}$ | |
| $\alpha_h(V) = 0.128 \exp(-(V + 50)/18)$ | |
| $\beta_h(V) = \frac{4}{1 + \exp(-(V + 27)/5)}$ | |
| $\alpha_n(V) = \frac{0.032(V + 52)}{1 - \exp(-(V + 52)/5)}$ | |
| $\beta_n(V) = 0.5 \exp(-(V + 57)/40)$ | |

In Eqs. (1)-(3), the letters *C*, *V*, *t* and *τ*, *g*, and *I* denote capacitance density, voltage, time, conductance density, and current density, respectively. The units that we use for these quantities are µF/cm^2^, mV, ms, mS/cm^2^, and µA/cm^2^. For brevity, units will usually be omitted hereafter. The parameter values of the model are *C* = 1, *gNa* = 100, *gK* = 80, *gL* = 0.1, *VNa* = 50, *VK* = -100, and *VL* = -67.

*I cells*: For fast-spiking PV+ interneurons, we use the model described in Wang and Buzsáki 1996^28^ . The Eqs. (1)-(4) are the same as in the E cell model, but the Eq. (5) is replaced by:

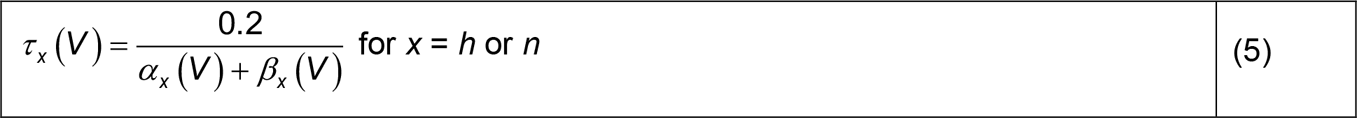

with the rate functions *αx* and *βx* (*x* = *m*, *h*, and *n*) are defined as follows:

|  |  |
| --- | --- |
| $\alpha_m(V) = \frac{0.1(V + 35)}{1 - \exp(-(V + 35)/10)}$ | |
| $\beta_m(V) = 4 \exp(-(V + 60)/18)$ | |
| $\alpha_h(V) = 0.07 \exp(-(V + 58)/20)$ | |
| $\beta_h(V) = \frac{1}{\exp(-0.1(V + 28)) + 1}$ | |
| $\alpha_n(V) = \frac{0.01(V + 34)}{1 - \exp(-0.1(V + 34))}$ | |
| $\beta_n(V) = 0.125 \exp(-(V + 44)/80)$ | |

The parameter values, using the same units as for the pyramidal cell, are *C* = 1, *gNa* = 35, *gK* = 9, *gL* = 0.1, *VNa* = 55, *VK* = −90, and *VL* = −65.

*O cells*: For the oriens lacunosum-moleculare (O-LM) interneurons, we use the model described in Tort et al. 2007^20^, which is a reduction of the multi-compartmental model described in Saraga et al. 2003^29^. The current-balance equation is given by:

|  |  |
| --- | --- |
| $C \frac{dV}{dt} = g_{Na} m^3 h (V_{Na} - V) + g_K n^4 (V_K - V) + g_A a b (V_A - V) + g_h r (V_h - V) + g_L (V_L - V) + I$ | (6) |

With

|  |  |
| --- | --- |
| $\frac{dx}{dt} = \frac{x_\infty(V) - x}{\tau_x(V)} \text{ for } x = m, h, n, a, b, r$ | (7) |

For *x* = *m*, *n*, *h*, the functions *x*_∞_(*V*) and *τ _x_* (*V*) are the same as in Eqs. (4) and (5), and the rate functions *αx* and *βx* are defined as follows:

|  |  |
| --- | --- |
| $\alpha_m(V) = \frac{-0.1(V + 38)}{\exp(-(V + 38)/10) - 1}$ | |
| $\beta_m(V) = 4 \exp(-(V + 65)/18)$ | |
| $\alpha_h(V) = 0.07 \exp(-(V + 63)/20)$ | |
| $\beta_h(V) = \frac{1}{1 + \exp(-(V + 33)/10)}$ | |
| $\alpha_n(V) = \frac{0.018(V - 25)}{1 - \exp(-(V - 25)/25)}$ | |
| $\beta_n(V) = \frac{0.0036(V - 35)}{\exp((V - 35)/12) - 1}$ | |

For *x* = *a*, *b*, *r*, the functions *x*_∞_(*V*) and *τ _x_* (*V*) are defined as:

For $x = a, b, r$ , the functions $x_\infty(V)$ and $\tau_x(V)$ are defined as:
|  |  |
| --- | --- |
| $a_\infty(V) = \frac{1}{1 + \exp(-(V + 14)/16.6)}$ | |
| $\tau_a(V) = 5$ | |
| $b_\infty(V) = \frac{1}{1 + 0.07 \exp((V + 71)/7.3)}$ | |
| $\tau_b(V) = \frac{1}{\frac{0.000009}{\exp((V - 26)/18.5)} + \frac{0.014}{0.2 + \exp(-(V + 70)/11)}}$ | |
| $r_\infty(V) = \frac{1}{1 + \exp((V + 84)/10.2)}$ | |
| $\tau_r(V) = \frac{1}{\exp(-14.59 - 0.086V) + \exp(-1.87 - 0.0701V)}$ | |

The parameter values, using the same units as for the pyramidal cell, are *C* = 1.3, *gL* = 0.05, *gNa* = 30, *gK* = 23, *gA* = 16, *gh* = 8, *VNa* = 90, *VK* = −100, *VA* = −90, *Vh* = −32, *VL* = −70; see also ref.^18^.

### Synaptic model

Following previous studies^17, 18, 27^, we model each synaptic input as a current of the form *Isyn,XY = GXY/NX s(V − Vsyn)*, where X and Y denote the type of the pre and postsynaptic cell, respectively (i.e. X and Y ɛ {E, I, O}), *GXY* is the maximal synaptic conductance, *NX* is the number of *X* cells, *V* is the membrane potential of the postsynaptic cell, and *Vsyn* is the reversal potential. The variable *s* is a normalized double-exponential function characterized by rise (*τ _X_*_,*r*_) and decay (*τ _X_*_,*d*_) time constants. The synapses were implemented using the Exp2Syn() built-in function of NEURON. The IPSPs originating from the O cells synapses were slower than IPSPs originating from the I cells synapses; following these stuides, we used *τ_E_*_,*r*_ = 0.05 ms and *τ_E_*_,*d*_ = 5.3 ms for E synapses, *τ_I_*_,*r*_ = 0.07 ms and *τ_I_*_,*d*_ = 9.1 ms for I synapses, and *τ_I_*_,*r*_ = 0.2 ms and *τ_I_*_,*d*_ = 22 ms for O synapses. The reversal potential was set to 0 mV for E synapses and to -80 mV for I and O synapses. We used *GII* = 0.01, *GOI* = 0.15, *GIO* = 0.2, *GIE* = 0.1, *GOE* = 0.12, *GEI* = 0.05, *GEO* = 0.09. In our model, in addition to the synaptic inputs (*Isyn,XY*) from X populations within CA1, the cells receive also some external drive currents, e.g. from other hippocampus regions or stimulus. We model this drive as having a constant (i.e. mean) level of *ε _X_* with an additive white noise. Here, we set *ε_E_* = 2, *ε_I_* = 0.3, and *ε_O_* = 0 . Here, for the E and I cells, *ε_E_* and *ε_I_* represent the mean excitatory drive currents to E and I populations through SC pathway, hence, hereafter, we denote them by *ε_SC_*_,*E*_ and *ε_SC_*_,*I*_, respectively. Using these drive currents, which are in consistent with previous studies^17, 18, 20^, simLFPs of Control model presented theta and gamma peak frequencies at ∼8-10 Hz and ∼48-50 Hz, respectively. Of note, the model gamma peak frequency provides a compromise between those we observed in HFS- and CCH-induced gamma oscillations (see Results)^30, 31^. The model theta peak frequency also closely reproduces that in our data (see Results).

### Model Local Field Potential

The simulated local field potential (simLFP) of the model consisted of the membrane potential of a “passive” E cell programmed inside the network^17, 18, 20^. This cell receives the same synaptic inputs as the “active” E cell. However, this cell does not send any synaptic output onto other cells and, in addition, it also does not spike given that its external drive current is set to zero. Power spectra of the simLFP was obtained using the Welch’s method (MATLAB *pwelch()* function) with 0.2 s Hamming windows, 75% overlap. In the preliminary analysis, we found that our findings hold true also for larger window sizes. However, our chosen window size was able to provide a clear visualization of the main results, by diminishing the potential harmonics of the theta and gamma rhythms. Accordingly, to show the average power spectrums and to read out the peak power robustly from simLFPs we used a window of 0.2 s size, and to extract the information about the exact peak frequency of the simLFPs we used a window of 2 s size (see Results); see also ref.^20^.

### Numeric and Random Aspects

All simulations were carried out using NEURON simulation program version 7.7 (https://www.neuron.yale.edu/neuron/)^32^. The model is publicly available code in ModelDB (https://senselab.med.yale.edu/ModelDB; accession number: 138421), and was simulated with a time step of 25 µs. As initial conditions, the membrane potential of each cell was uniformly distributed between -85 and -60 mV and the channel-gating variables were set to their corresponding steady-state values. Each cell was further randomized using a clamping current (*IClamp*) of random uniform magnitude and random uniform duration between 0 and *tsyn*/2, where *tsyn* = 0.5 s is the time when synapses were turned on. In addition to the drive current (see previous section), each cell in the network also received additive white noise current inputs (see also^20, 21^). For each of the Control and NMDAR-Ab models (see Supplementary Table 2), we simulated the model for 50 trials of 12 s and used the last 10 s of simLFPs for the analysis. Note that, in simLFP signals, the mentioned randomizations, together with the stochastic components of the drive currents, varied the exact theta and gamma peak frequencies as well as their harmonics from trial to trial thereby yielding the broadening of the peak frequencies by ∼2 Hz in the trial-averaged spectrum results (see Results).

### Modulation of network oscillations

Our model revealed an increase in θ- and to a lesser degree in γ-oscillations after implementing the NMDAR-induced reduction in *ε_SC_*_,*E*_ individually (see model 3 in Fig. 7d). To evaluate the potential compensatory role of the reduction in decay time constants of synaptic inhibition mediated by O-cells (*τ_O_*_,*d*_) and I-cells (*τ_I_*_,*d*_) we computed the power modulation parameter (Fig. 7f):

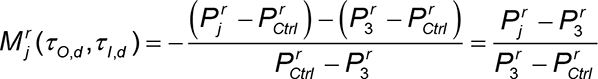

Where, *r* ∈{θ,γ},*^r^* is the peak power of r-type oscillations (*r* ∈{θ,γ}) in model *j*, where *j* refers to using different set of reductions (0, 30, 60, and 80%) in *τ_O_*_,*d*_ and *τ_I_*_,*d*_ in NDMAR-Ab basal model relative to the Control model. are related to NMDAR-Ab model #3 and Control model (Fig. 7d), respectively. Intuitively, *^r^* quantifies the amount change in excessive θ- and γ-oscillatory powers arising from the reduced *ε_SC_*_,*E*_ . Note that the denominator of *^r^* is positive for both θ- and γ-oscillations. By this definition, *M^r^*= −1 represents a full (i.e. 100%) suppression of the excessive power, bringing it back to the Control-model level. A *^r^* with values of 0 and >0 indicte no modulation and an increase of (i.e. deliberating) the excessive power, respectively. Peak powers were computed based on the power spectrums in the corresponding frequency bands.

### Statistical analysis

Statistical analyses were performed using OriginPro 2019, Matlab 2020a, and SPSS 27. All data are reported as mean ± standard error of the mean (SEM), if not stated otherwise. The Shapiro–Wilk test was used to test for normality. The *F*-test was used to test for homogeneity of variances. Parametric testing procedures were applied for normally distributed data; otherwise, nonparametric tests were used. Except for the Shapiro–Wilk test and *F*-test where P values < 0.05 were considered statistically significant, the actual P values were stated for other tests. Bin-wise permutation tests in Fig. 6e,i and Supplementary figure 5) were performed at a significance level of 5% using Two-tailed permutation test of Cohen thereby accounting for multiple comparison problem (see Methods for detail); the significant bins were designated by a star symbol. Whole-curve permutation (Figs. 2b,f,g,h, and 4f, and Supplementary figure 3a) tests were performed by computing the area-under-curve (AUC) of the group-averaged curve, followed by computing the difference of AUCs of the two groups (empirical-difference; eDiff). The individual curves were then shuffled across the groups and the difference was re-calculated (shuffled-difference; sDiff). This was repeated for two million times. Finally, the P value (two-tailed) for eDiff was computed as the number of times that sDiff was bigger than |eDiff| or smallaer than -|eDiff|, divided by the number of shuffling times. Details of the applied statistical tests with the sample sizes are provided in Tables Supplementary Table 1.

## Supplementary Figures

**Supplementary figure 1.**
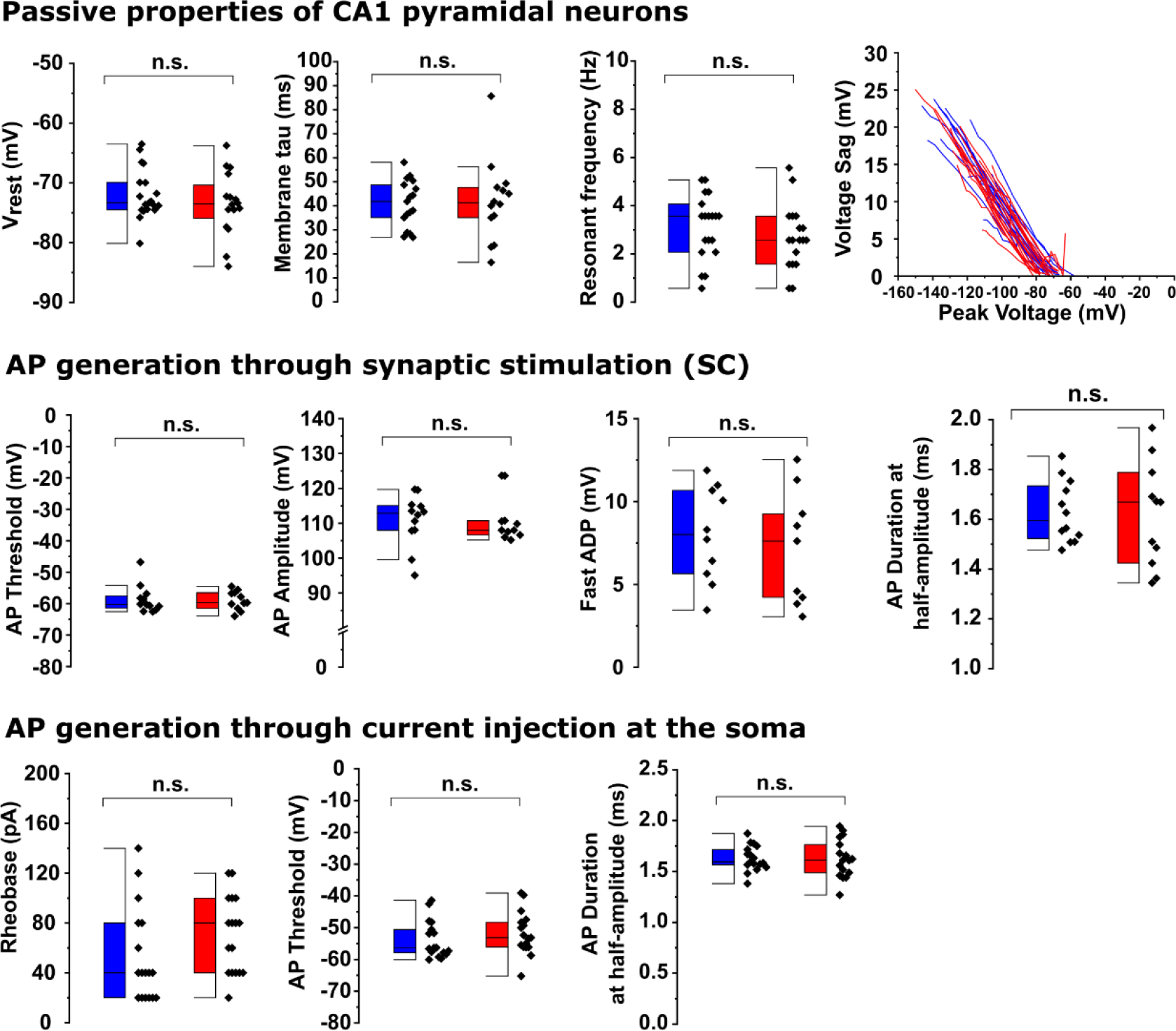
NMDAR-Ab do not change action potential properties of CA1 pyramidal neurons. *Upper row:* Additional passive properties of CA1-PNs. The resting membrane potential, the membrane time constant, the resonant frequency as well as the voltage sag were not changed following NMDAR-Ab treatment. *Middle row:* Action potentials (APs) of CA1-PNs were elicited by single-pulse stimulation of afferent Schaffer collaterals (SC). We found no change in the AP threshold, AP amplitude, fast afterdepolarization (ADP) or AP duration at half-amplitude. *Lower row:* APs were elicited by a constant current-step injection at the soma for 1 second. We found no change in the rheobase, AP threshold or AP duration at half-amplitude. See Supplementary Table 1 for statistical details.

**Supplementary figure 2.**
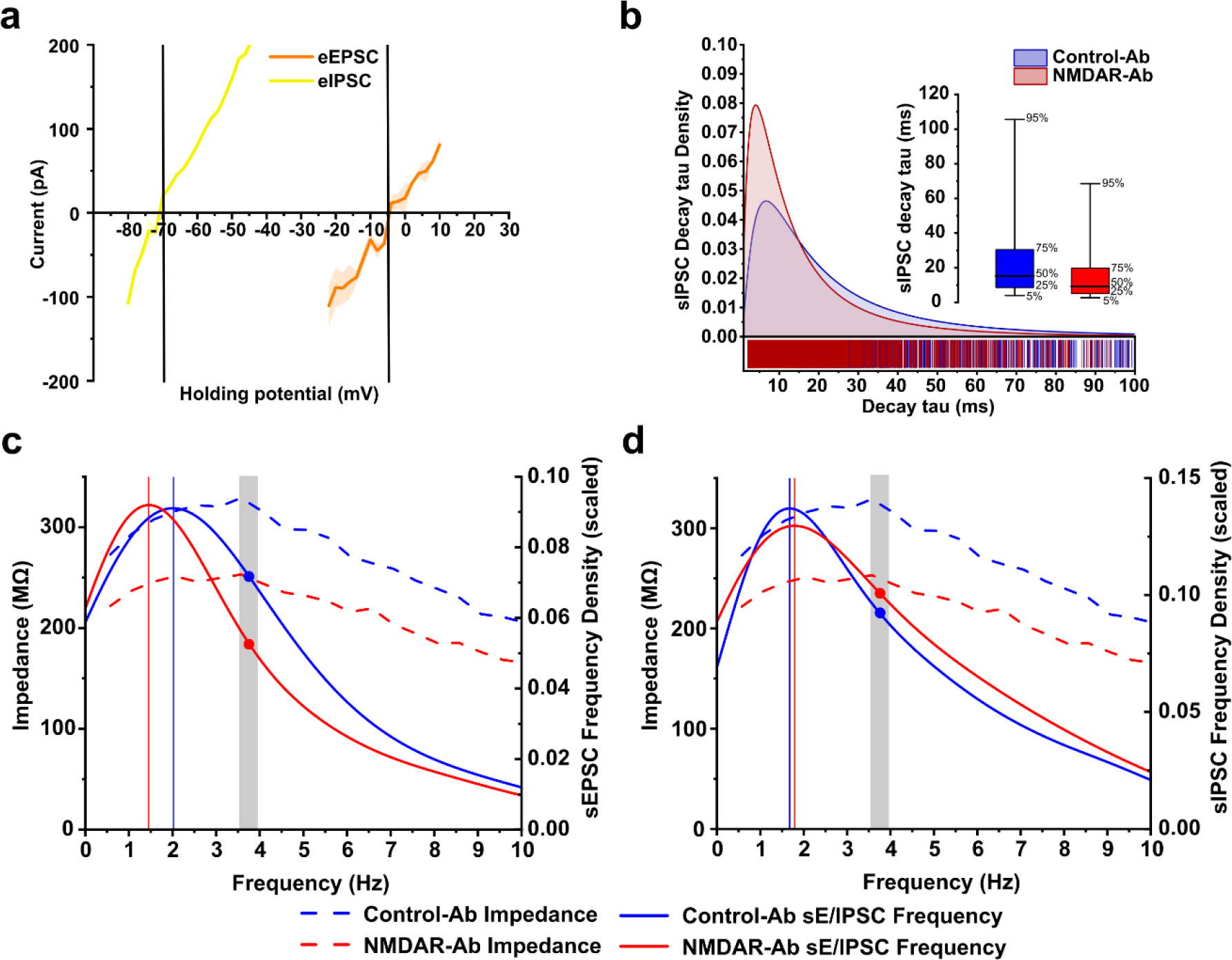
NMDAR-Ab diverges sEPSC dominant frequency from CA1- PN membrane resonant frequency. **(a)** The reversal potentials of synaptic excitation (with 100µM PTX in the bath solution) and inhibition (with 10µM CNQX and 50mM AP-5 in the bath solution) at CA1 PNs were analyzed in untreated hippocampal slices. The experimentally determined reversal potentials were -5 mV and -70 mV (vertical lines), respectively. **(b)** The density distribution of sIPSC decay tau was shifted toward lower values for both fast and slow (inset) decays. **(c)** The dominant frequency (vertical line) of sEPSC (solid line), and the intrinsic membrane resonant frequency of CA1 PNs diverged after NMDAR-Ab treatment, while the density of sEPSC events was reduced at the peak resonant frequency (spatial distance between the intersections of grey line with the sEPSC frequency distributions), unveiling a further mechanism underlying the impaired excitatory drive. **(d)** Same as (c), but for sIPSC. There was no apparent change in the overlap of sIPSC dominant frequency and the resonance of PN after NMDAR-Ab treatment.

**Supplementary figure 3.**
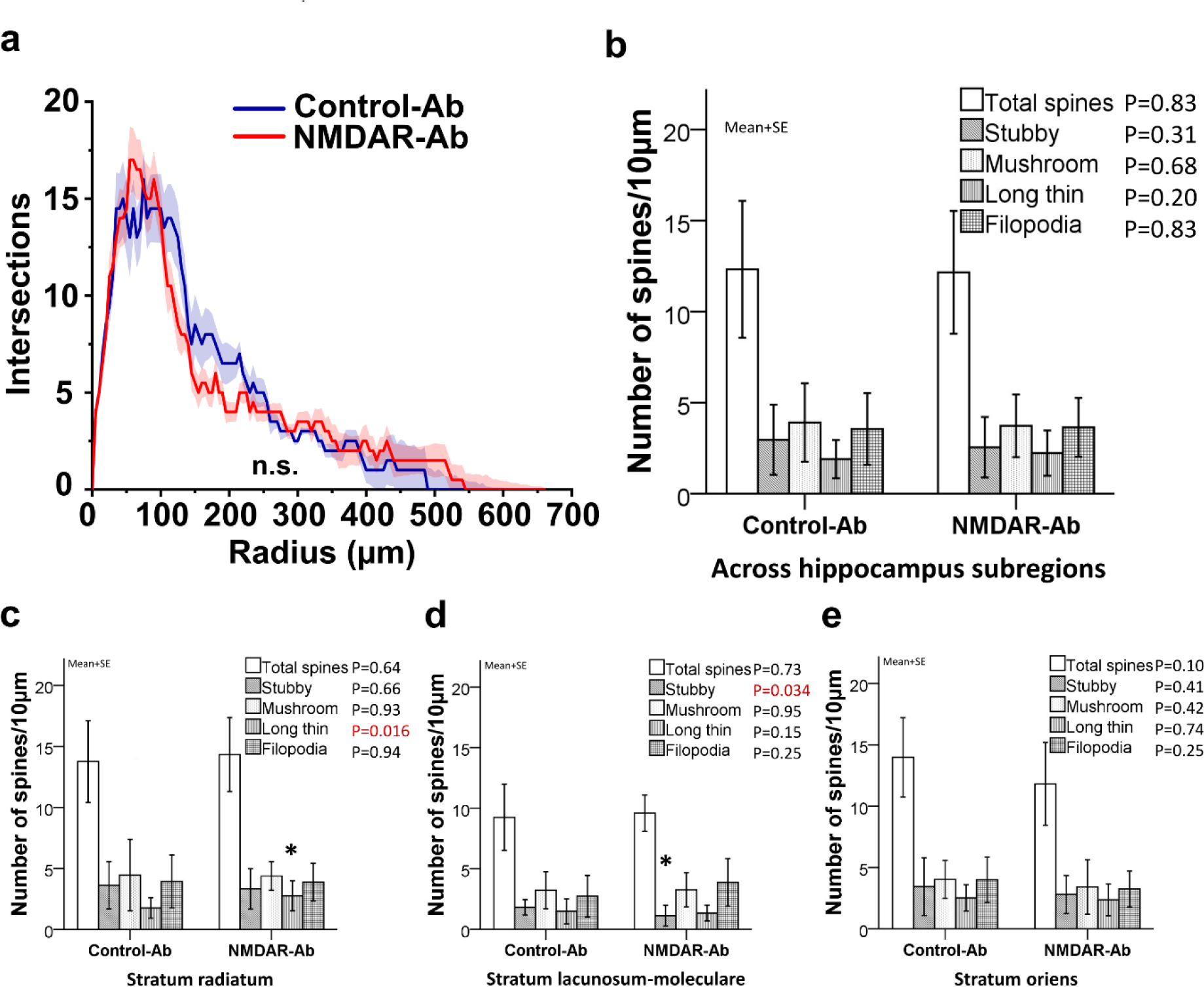
NMDAR-Ab causes no relevant changes in the dendritic morphology of CA1 PNs. **(a)** Sholl analysis of dendritic trees revealed no change in the morphology of CA1-PNs. The mean (solid line) ± SEM (shaded area). **(b)** Reconstruction of Biocytin-filled CA1-PNs did not show a significant change in the overall number or morphology of dendritic spines across the treatments. **(c,d,e)** Detailed analysis of regional spine distributions revealed only minor changes in the number of long thin spines and stubby spines in the strata radiatum and lacunosum-moleculare of CA1, respectively.

**Supplementary figure 4.**
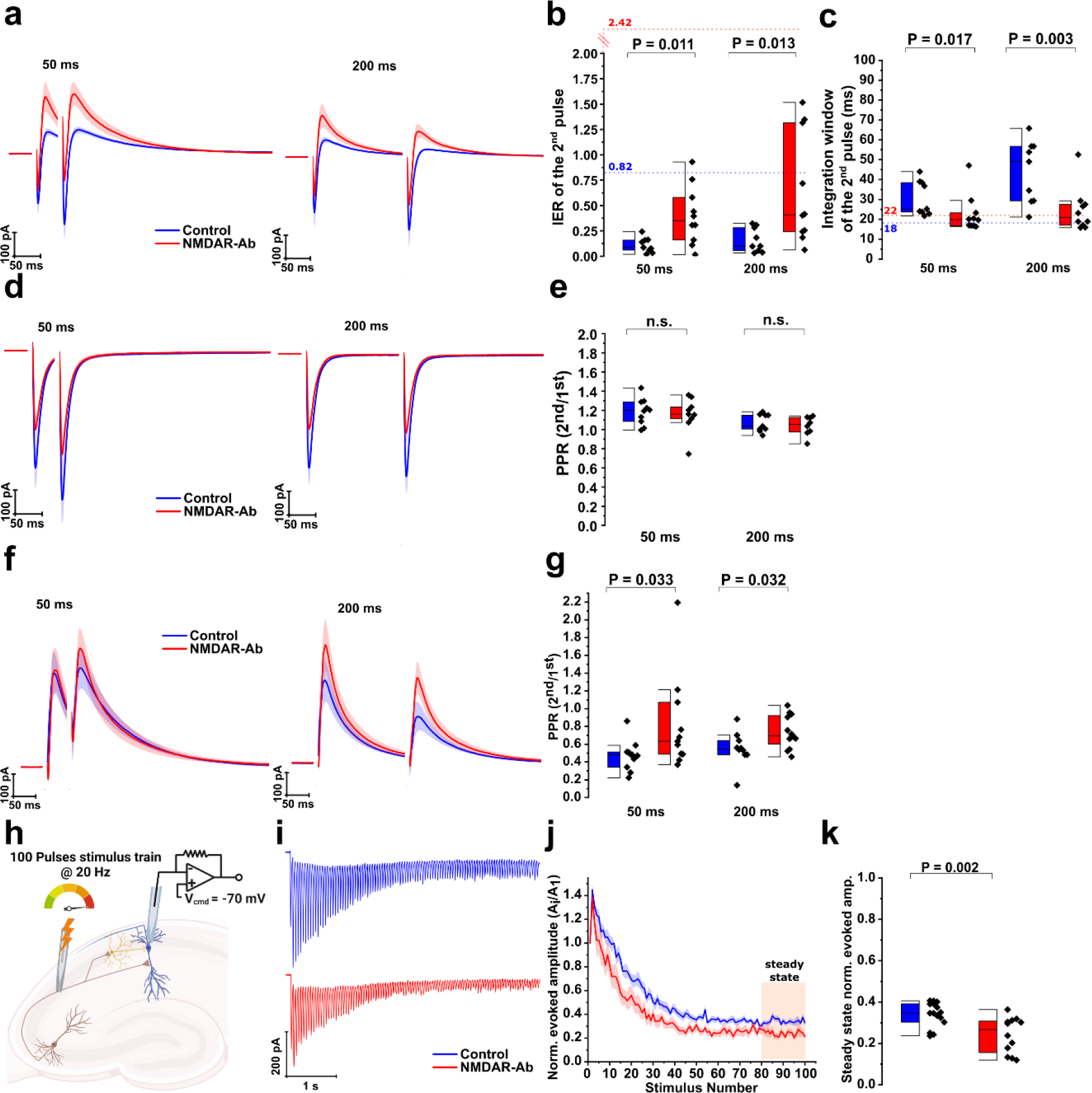
NMDAR-Ab induces cell-type specific changes in short-term plasticity (STP) within the hippocampal feed-forward (FF) network. Schaffer collaterals (SC) received paired-pulse stimulations (PPS) at 50ms or 200ms and CA3→CA1 responses of the FF network were recorded at different holding potentials. **(a)** Average biphasic responses (at a holding potential of ca. -35mV) onto PNs after PPS delivered at *(left panel)* 50ms or *(right panel)* 200ms intervals. **(b)** During STP, the inhibitory-excitatory ratio (IER) of the second pulse had higher values under NMDAR-Ab than in Control-Ab group, at both stimulation intervals. However, in both groups, the IER of the second pulse was lower than of the first pulse (depicted by dashed lines together with the corresponding values adopted from Fig. 4). **(c)** During STP, the integration window of the second pulse was shorter for NMDAR- Ab group as compared to Control-Ab values, where the integration window increased relative to the first pulse (dashed lines together with the corresponding values in ms adopted from Fig. 4). **(d)** Same as in (a) but for eEPSC (at a holding potential of -70mV). **(e)** We found no change in the paired-pulse ration (PPR) of monosynaptic excitation after NMDAR-Ab treatment. **(f)** Same as in (a) but for bisynaptic eIPSC (at a holding potential of -5mV). **(g)** There was less short-term depression of bisynaptic eIPSC following NMDAR-Ab treatment at both stimulation intervals. **(h-k)** CA1-PN received 100 pulses at 20 Hz at maximal stimulation intensity to investigate excitatory transmission during AP trains (at a holding potential of -70mV). **(h)** Schematic of the recording protocol. **(i)** Averaged eEPSC trains in response to 20 Hz stimulation train. **(j)** Evoked EPSCs, normalized to the first amplitude, exhibited a time- dependent decrease in amplitude before reaching a steady state (last 20 stimuli). Note the more pronounced, progressive decrease in the amplitude after NMDAR-Ab treatment. The mean (solid line) ± SEM (shaded area). **(k)** NMDAR-Ab led to a drop in the steady-state (last 20 stimuli) level of normalized eEPSCs amplitudes. The steady state duration was designated by the pink-colored shaded area in (j).

**Supplementary figure 5.**
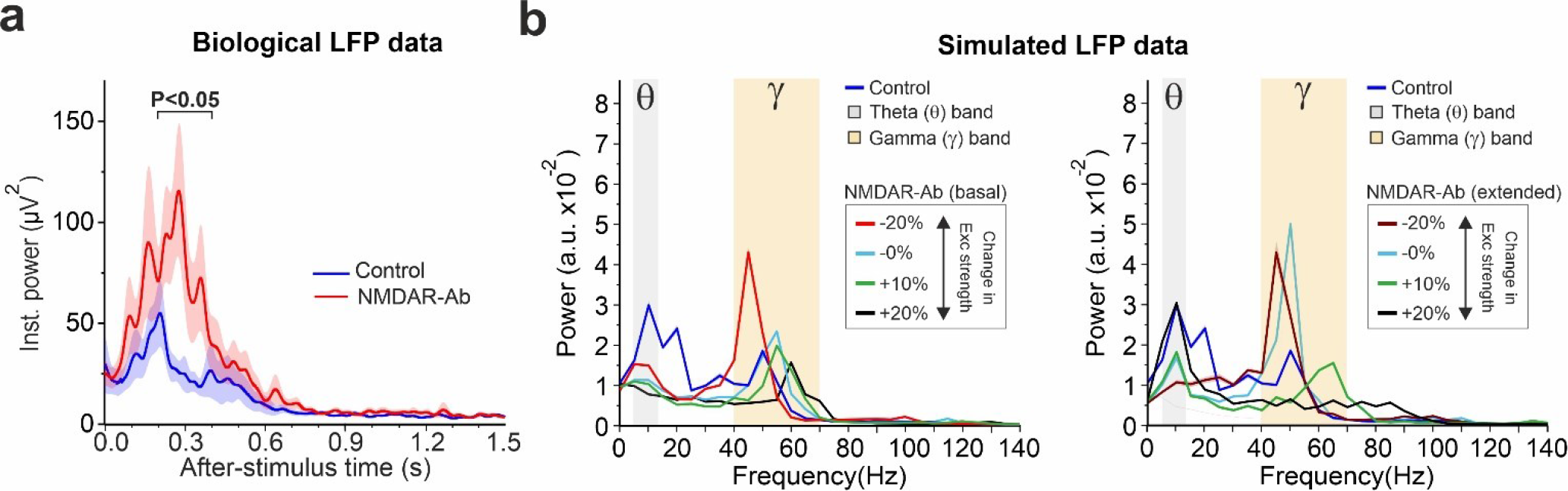
Additional analysis related to Figures 6 and 7. **(a)** The amplification of γ-oscillations under NMDAR-Ab is also present in the raw (i.e. non- normalized) instantaneous power traces of HFS-induced LFPs. Same format is used as in Fig. 6e. **(b)** Augmentation of Exc synaptic strength in NMDAR-Ab basal and extended models suppresses the excessive γ-oscillatory power. This manipulation in the latter model tends to also boost the reduced θ-oscillatory power. Same format is used as in Fig. 8d. The changes in Exc synaptic strengths are relative to that in the Control model. In NMDAR-Ab basal model these modifications are applied only to *ε_SC_*_,*E*_, but in NMDAR-Ab extended model they are applied to all Exc synaptic connections within CA1, namely to *ε_SC_*_,*E*_, *ε_SC_*_,*I*_, *G_EI_*, and *^G^EO*.

## Supplementary Tables

**Table S1.** Synopsis of statistical tests.

| Related to | Descriptive statistics | Test(s) | Test statistics |
| --- | --- | --- | --- |
| **Figure 1** | *Group: mean ± SEM (no. of cells/no. of mice)* |  |  |
| 1e |  |  |  |
| I/O Curves | Control-Ab: 12/7<br>NMDAR-Ab: 11/5 |  |  |
| 1f |  |  |  |
| I/O threshold | Control-Ab: 2.61438 ± 0.20293 (12/7)<br>NMDAR-Ab: 4.07991 ± 0.51848 events (11/5) | Two-sample t-test | P = 0.02068<br>t = -2.63214<br>df = 13.0207 |
| 1g |  |  |  |
| I/O gain | Control-Ab: 0.31153 ± 0.02198 (12/7)<br>NMDAR-Ab: 0.33223 ± 0.077 (11/5) | Two-sample t-test | P = 0.80054<br>t = -0.2585<br>df = 11.626 |
| 1h |  |  |  |
| Lower panel EPSP<br>amp./EPSP slope | Control-Ab: 612 events (13/7)<br>NMDAR-Ab: 642 events (10/5) | Linear mixed<br>model | β = 0.28<br>t = 2.27<br>P = 0.023 |
| 1k |  |  |  |
| EPSP jitter | Control-Ab: 2.19534 ± 0.36297 (12/7)<br>NMDAR-Ab: 0.95792 ± 0.28894 (11/5) | Mann-Whitney<br>test | P = 0.0225<br>Z = 2.24642<br>U = 103 |
| 1l |  |  |  |
| AP jitter | Control-Ab: 1.85938 ± 0.38191 (12/7)<br>NMDAR-Ab: 0.93912 ± 0.17858 (11/5) | Mann-Whitney<br>test | P = 0.0268<br>Z = 2.18487<br>U = 102 |
| **Supplementary**<br>**Fig. 1 – Upper row** | *Group: mean ± SEM (no. of cells/no. of mice)* |  |  |
| V<sub>rest</sub> | Control-Ab: -71.9217 ± 0.9567 (19/7)<br>NMDAR-Ab: -73.4874 ± 1.3291 (16/6) | Two-sample t-test | P = 0.33596<br>t = 0.97641<br>df = 33 |
| Membrane tau | Control-Ab: 40.9805 ± 2.36247 (17/7)<br>NMDAR-Ab: 41.90945 ± 4.49286 (14/6) | Two-sample t-test | P = 0.85664<br>t = -0.183<br>df = 33 |
| Resonant frequency | Control-Ab: 3.12403 ± 0.30564 (17/7)<br>NMDAR-Ab: 2.68251 ± 0.31195 (17/6) | Two-sample t-test | P = 0.31905<br>t = 1.0108<br>df = 35 |
| **Supplementary**<br>**Fig. 1 – Middle panel** |  |  |  |
| AP threshold | Control-Ab: -58.635 ± 1.2844 (12/7)<br>NMDAR-Ab: -58.9997 ± 0.9095 (11/5) | Mann-Whitney<br>test | P = 0.6947<br>Z = -0.40005<br>U = 59 |
| AP amplitude | Control-Ab: 110.8137 ± 2.13 (12/7)<br>NMDAR-Ab: 110.8947 ± 1.975 (11/5) | Mann-Whitney<br>test | P = 0.37934<br>Z = 0.89241<br>U = 19.94385 |
| Fast ADP | Control-Ab: 8.01388 ± 0.90259 (10/7)<br>NMDAR-Ab: 7.214 ± 1.15322 (9/5) | Two-sample t-test | P = 0.58812<br>t = 0.55202<br>df = 17 |
| AP Duration at half-amplitude | Control-Ab: 1.62876 ± 0.03587 (12/7)<br>NMDAR-Ab: 1.61765 ± 0.06295 (11/5) | Two-sample t-test | P = 0.87693<br>t = 0.15676<br>df = 21 |
| **Supplementary**<br>**Fig. 1 – Lower panel** |  |  |  |
| Rheobase | Control-Ab: 52.941 ± 9.226 (17/6)<br>NMDAR-Ab: 70.588 ± 7.499 (17/6) | Mann-Whitney<br>test | P = 0.07753<br>Z = -1.7594<br>U = 94 |
| AP threshold | Control-Ab: -53.747 ± 1.4261 (17/6)<br>NMDAR-Ab: -51.8004 ± 1.617 (17/6) | Mann-Whitney<br>test | P = 0.15983<br>Z = -1.41218<br>U = 103 |
| AP Duration at half-amplitude | Control-Ab: 1.62809 ± 0.03011 (17/6)<br>NMDAR-Ab: 1.62737 ± 0.04543 (17/6) | Two-sample t-test | P = 0.98952<br>t = 0.01324<br>df = 32 |
| **Figure 2** |  |  |  |
| 2b |  |  |  |
| ZAP | Control-Ab: (19/7) For frequencies up to<br>NMDAR-Ab: (18/6) alpha (12 Hz) | Two-tailed whole-curve permutation test, 2 million shuffles | P = 0.0471 |
| 2c |  |  |  |
| Maximal impedance | Control-Ab: 350.62 ± 28.48 (19/7)<br>NMDAR-Ab: 273.82 ± 19.96 (18/6) | Two-sample t-test | P = 0.03557<br>t = 2.18627<br>df = 35 |
| 2d |  |  |  |
| Input resistance | Control-Ab: 262.35712 ± 19.32231 (17/7)<br>NMDAR-Ab: 204.86977 ± 17.38323 (14/6) | Two-sample t-test | P = 0.03851<br>t = 2.16798<br>df = 29 |
| 2f |  |  |  |
| Average inst. frequency | Control-Ab: (17/7)<br>NMDAR-Ab: (17/6) | Two-tailed whole-curve permutation test, 2 million shuffles | P = 0.0036 |
| 2g |  |  |  |
| Number of Aps/1s step | Control-Ab: (17/7)<br>NMDAR-Ab: (17/6) | Two-tailed whole-curve permutation test, 2 million shuffles | P = 0.3241 |
| 2h |  |  |  |
| SFA | Control-Ab: (17/7)<br>NMDAR-Ab: (17/6) | Two-tailed whole-curve permutation test, 2 million shuffles | P = 0.44895 |
| 2i |  |  |  |
| Cells with rebound APs | Control-Ab: 6/18 (18/7)<br>NMDAR-Ab: 1/17 (17/6) | Chi-square | $\chi^2$ (1, N = 35) = 4.1176<br>P = 0.042438 |
| **Supplementary Fig. 2** |  |  |  |
| SF2a | eEPSC: (5/1)<br>eIPSC: (4/1) |  |  |
| SF2b | Control-Ab: 2594 events/12 cells<br>NMDAR-Ab: 2811 events/14 cells |  |  |
| SF2c | Control-Ab: ZAP: 19/7; sEPSC: 12/6<br>NMDAR-Ab: ZAP: 18/6; sEPSC: 14/6 |  |  |
| SF2d | Control-Ab: ZAP: 19/7; sIPSC: 7/3<br>NMDAR-Ab: ZAP: 18/6; sIPSC: 8/3 |  |  |
| **Figure 3** |  |  |  |
| 3d |  |  |  |
| sEPSC amplitude | Control-Ab: 18.6765 ± 1.4999 (12/6)<br>NMDAR-Ab: 14.63475 ± 1.0622 (14/6) | Two-sample t-test | P = 0.0342<br>t = 2.24575<br>df = 24 |
| 3e |  |  |  |
| sEPSC frequency | Control-Ab: 1.82151 ± 0.36212 (12/6)<br>NMDAR-Ab: 1.42697 ± 0.3051 (14/6) | Two-sample t-test | P = 0.40946<br>t = 0.83952<br>df = 24 |
| 3f |  |  |  |
| sEPSC decay tau | Control-Ab: 5.7075 ± 0.40908 (12/6)<br>NMDAR-Ab: 4.39801 ± 0.27474 (14/6) | Two-sample t-test | P = 0.01183<br>t = 2.72436<br>df = 24 |
| 3g |  |  |  |
| sEPSC charge transfer | Control-Ab: 171.2298 ± 18.133 (12/6)<br>NMDAR-Ab: 117.5393 ± 9.958 (14/6) | Two-sample t-test | P = 0.01259<br>t = 2.69717<br>df = 24 |
| 3k |  |  |  |
| sIPSC amplitude | Control-Ab: 22.45 ± 1.6761 (12/6)<br>NMDAR-Ab: 18.8784 ± 1.9595 (14/6) | Two-sample t-test | P = 0.19587<br>t = 1.3635<br>df = 13 |
| 3l |  |  |  |
| sIPSC frequency | Control-Ab: 2.1981 ± 0.59024 (12/6)<br>NMDAR-Ab: 2.39497 ± 0.6669 (14/6) | Mann-Whitney test | P = 0.95509<br>Z = -0.05786 |
|  |  |  | U = 27 |
| 3m |  |  |  |
| sIPSC decay tau | Control-Ab: 15.6584 ± 2.14365 (12/6)<br>NMDAR-Ab: 7.0362 ± 0.916 (14/6) | Two-sample t-test | P = 0.00585<br>t = 3.69873<br>df = 8.15755 |
| 3n |  |  |  |
| sIPSC charge transfer | Control-Ab: 689.4082 ± 48.74 (12/6)<br>NMDAR-Ab: 362.9173 ± 40.708 (14/6) | Two-sample t-test | P = 1.7619E-4<br>t = 5.18312<br>df = 13 |
| **Supplementary Fig. 3** |  |  |  |
| SF3a. Sholl analysis | Control-Ab: (10/8)<br>NMDAR-Ab: (12/4) | Two-tailed whole-curve permutation test, 2 million shuffles | P = 0.95199 |
| SF3b-e. | Control-Ab: (10/8)<br>NMDAR-Ab: (12/4) | ANOVA | See figure |
| **Figure 4** |  |  |  |
| 4c |  |  |  |
| IER | Control-Ab: 0.82384 ± 0.17027 (10/4)<br>NMDAR-Ab: 2.42871 ± 0.65684 (10/3) | Two-sample t-test | P = 0.03913<br>t = -2.36514<br>df = 10.20412 |
| 4d |  |  |  |
| Integration window | Control-Ab: 22.5642 ± 1.808 (10/4)<br>NMDAR-Ab: 18.0008 ± 1.792 (10/3) | Two-sample t-test | P = 0.08984<br>t = 1.7927<br>df = 18 |
| 4f |  |  |  |
| eEPSC I/O curves | Control-Ab: (12/6)<br>NMDAR-Ab: (14/6) | Two-tailed whole-curve permutation test, 2 million shuffles | P = 0.000437 |
| 4g |  |  |  |
| eEPSC max. amplitude | Control-Ab: 426.7183 ± 41.416 (20/5)<br>NMDAR-Ab: 266.5814 ± 25.817 (22/4) | Two-sample t-test | P = 0.00179<br>t = 3.34609<br>df = 40 |
| 4h |  |  |  |
| eEPSC decay tau | Control-Ab: 20.9179 ± 2.1461 (20/5)<br>NMDAR-Ab: 19.05976 ± 2.0968 (22/4) | Two-sample t-test | P = 0.54011<br>t = 0.61747<br>df = 40 |
| 4k |  |  |  |
| eIPSC amplitude | Control-Ab: 369.5357 ± 41.172 (11/4)<br>NMDAR-Ab: 625.035 ± 116.315 (11/3) | Two-sample t-test | P = 0.05975<br>t = -2.07072<br>df = 12.46713 |
| 4l |  |  |  |
| eIPSC decay tau | Control-Ab: 73.45614 ± 5.9271 (11/4)<br>NMDAR-Ab: 46.69633 ± 4.318 (11/3) | Two-sample t-test | P = 0.0016<br>t = 3.64916<br>df = 20 |
| **Supplementary Fig. 4** |  |  |  |
| SF4b. - IER PP 50ms | Control-Ab: 0.10885 ± 0.02411 (9/4)<br>NMDAR-Ab: 0.4005 ± 0.09131 (10/3) | Two-sample t-test | P = 0.01117<br>t = -3.08807<br>df = 10.2428 |
| SF4b. - IER PP 200ms | Control-Ab: 0.15738 ± 0.03976 (9/4)<br>NMDAR-Ab: 0.64629 ± 0.17283 (10/3) | Mann-Whitney test | P = 0.01327<br>Z = -2.40866<br>U = 15 |
| SF4c. - IW PP 50ms | Control-Ab: 30.63778 ± 2.90645 (9/4)<br>NMDAR-Ab: 22.70917 ± 2.98683 (10/3) | Mann-Whitney test | P = 0.01721<br>Z = 2.32702<br>U = 74 |
| SF4c. - IW PP 200ms | Control-Ab: 44.01107 ± 5.23668 (9/4)<br>NMDAR-Ab: 24.56275 ± 3.48788 (10/3) | Mann-Whitney test | P = 0.00299<br>Z = 2.81691<br>U = 80 |
| SF4e. - PPR 50ms | Control-Ab: 1.18649 ± 0.04297 (10/4)<br>NMDAR-Ab: 1.15367 ± 0.06023 (9/3) | Two-sample t-test | P = 0.65805<br>t = 0.45049<br>df = 17 |
| SF4e. - PPR 200ms | Control-Ab: 1.06812 ± 0.03067 (9/4)<br>NMDAR-Ab: 1.03784 ± 0.03546 (8/3) | Two-sample t-test | P = 0.52599<br>t = 0.64925<br>df = 15 |
| SF4g. - PPR 50ms | Control-Ab: $0.47184 \pm 0.05087$ (11/4)<br>NMDAR-Ab: $0.76029 \pm 0.15409$ (11/3) | Mann-Whitney test | $P = 0.0336$<br>$Z = -2.10128$<br>$U = 28$ |
| SF4g. - PPR 200ms | Control-Ab: $0.5515 \pm 0.05972$ (10/4)<br>NMDAR-Ab: $0.7368 \pm 0.05385$ (12/3) | Mann-Whitney test | $P = 0.03188$<br>$t = -2.30695$<br>$df = 20$ |
| SF4k. | Control-Ab: $0.34027 \pm 0.01527$ (15/5)<br>NMDAR-Ab: $0.24127 \pm 0.02483$ (12/4) | Two-sample t-test | $P = 0.0016$<br>$t = 3.54036$<br>$df = 25$ |
| **Figure 5** | Proteomics |  |  |
|  | see Supplementary Online Data |  |  |
| **Figure 6** | *Group: mean ± SEM (no. of slices)* |  |  |
| 6e |  |  |  |
| Instantaneous power | Control-Ab (n = 8)<br>NMDAR-Ab (n = 9) | Two-tailed permutation test of Cohen, 1 million shuffles | $P < 0.05$ |
| 6f |  |  |  |
| Peak frequency | Control-Ab: $70.1875 \pm 4.55025$ (n = 8)<br>NMDAR-Ab: $75.7222 \pm 2.88809$ (n = 9) | Two-sample t-test | $P = 0.3097$<br>$t = -1.05154$<br>$df = 15$ |
| Bandwidth | Control-Ab: $26 \pm 2.5$ (n = 8)<br>NMDAR-Ab: $30.722 \pm 3.089$ (n = 9) | Two-sample t-test | $P = 0.2607$<br>$t = -1.1688$<br>$df = 15$ |
| 6g (inset) |  |  |  |
| Power | Control-Ab: $2.76E-6 \pm 5.16E-7$ (n = 8)<br>NMDAR-Ab: $1.55E-6 \pm 2.40E-7$ (n = 9) | Two-sample t-test | $P = 0.04328$<br>$t = 2.20738$<br>$df = 15$ |
| 6i |  |  |  |
| TGC | Control-Ab (n = 8)<br>NMDAR-Ab (n = 9) | Two-tailed permutation test of Cohen, 1 million shuffles | $P < 0.05$ |
| **Supplementary Fig. 5a** |  |  |  |
| Instantaneous power (non-normalized) | Control-Ab (n = 8)<br>NMDAR-Ab (n = 9) | Two-tailed permutation test of Cohen, 1 million shuffles | $P < 0.05$ |
| **Figure 8** |  |  |  |
| 8a |  |  |  |
| Peak power | Control-Ab (CCH): $1.07E-7 \pm 2.82E-8$ (n = 7)<br>NMDAR-Ab (CCH): $2.60E-7 \pm 1.59E-7$ (n = 10) | Two-sample t-test | $P = 0.03284$<br>$t = -2.3506$<br>$df = 15$ |
| Peak frequency | Control-Ab (CCH): $25.809 \pm 0.345$ (n = 7)<br>NMDAR-Ab (CCH): $25.634 \pm 0.257$ (n = 10) | Mann-Whitney U-test (exact) | $P = 0.78116$<br>$Z = 0.36438$<br>$U = 39$ |
| Results section of Figure 8 |  |  |  |
| Peak power | Control-Ab (CCH): $1.07E-7 \pm 2.82E-8$ (n = 7)<br>NMDAR-Ab (CCH+PAM): $2.52E-7 \pm 8.01E-8$ (n = 10) | Mann-Whitney U-test (exact) | $P = 0.36384$<br>$Z = -0.92711$<br>$U = 25$ |
| Figure 8c (inset) |  |  |  |
| Peak power | Control-Ab (CCH+PAM): $1.03E-7 \pm 2.88E-8$ (n = 7)<br>NMDAR-Ab (CCH+PAM): $2.52E-7 \pm 8.01E-8$ (n = 10) | Mann-Whitney U-test (exact) | $P = 0.41732$<br>$Z = -0.82952$<br>$U = 26$ |
| Results section of Figure 8 |  |  |  |
| Peak power | Control-Ab (CCH): $1.07E-7 \pm 2.82E-8$ (n = 7)<br>Control-Ab (CCH+PAM): $1.03E-7 \pm 2.87E-8$ (n = 7) | Paired-sample t-test | $P = 0.77456$<br>$t = 0.29964$<br>$df = 6$ |
| Figure 8d (inset) |  |  |  |
| 0.5-Lag | Control-Ab (CCH): $0.05136 \pm 0.00198$ (n = 7)<br>NMDAR-Ab (CCH): $0.06172 \pm 0.00406$ (n = 10) | Mann-Whitney U-test (exact) | $P = 0.03538$<br>$Z = -2.03712$<br>$U = 14$ |
| Figure 8e (inset) |  |  |  |
| 0.5-Lag | Control-Ab (CCH): $0,05136 \pm 0,00198$ (n = 7)<br>NMDAR-Ab (CCH+PAM): $0,05665 \pm 0,00274$ (n = 10) | Mann-Whitney U-test (exact) | P = 0.14685<br>Z = -1.42701<br>U = 20.5 |
| Results section of Figure 8 |  |  |  |
| 0.5-Lag | Control-Ab (CCH): $0,05136 \pm 0,00198$ (n = 7)<br>Control-Ab (CCH+PAM): $0,05246 \pm 0,00306$ (n = 7) | Wilcoxon signed-rank test (exact) | P = 0.79688<br>Z = 0.25446<br>W = 16 |

**Supplementary Table 2.** Parameter sets used in NMDAR-Ab network model.

|  | 1 | 2 | 3 | 4 | 5 | 6 | 7 | 8 |
| --- | --- | --- | --- | --- | --- | --- | --- | --- |
| $\varepsilon_{SC,E}$ | -20 | -20 | -20 | 0 | -20 | -20 | 0 | -20 |
| $\varepsilon_{SC,I}$ | 0 | 0 | 0 | 0 | 0 | 0 | 0 | -20 |
| $\tau_{I,d}$ | 0 | 0 | 0 | 0 | -60 | 0 | 0 | -30 |
| $\tau_{O,d}$ | -60 | X | 0 | -60 | 0 | -60 | 0 | -60 |
| $G_{EI}$ | 0 | 0 | 0 | 0 | 0 | 0 | 0 | -20 |
| $G_{EO}$ | 0 | 0 | 0 | 0 | 0 | 0 | 0 | -20 |
| $G_{OI}$ | 0 | X | 0 | 0 | 0 | 0 | 0 | 0 |
| $G_{OE}$ | 0 | X | 0 | 0 | 0 | 0 | 0 | 0 |
| $R_{in}$ | 0 | 0 | 0 | 0 | 0 | -20 | -20 | -20 |

